# Helminth exposure protects against murine SARS-CoV-2 infection through macrophage dependent T cell activation

**DOI:** 10.1101/2022.11.09.515832

**Authors:** Kerry L. Hilligan, Oyebola O. Oyesola, Sivaranjani Namasivayam, Nina Howard, Chad S. Clancy, Sandra D. Oland, Nicole L. Garza, Bernard A. P. Lafont, Reed F. Johnson, Katrin D. Mayer-Barber, Alan Sher, P’ng Loke

## Abstract

Helminth endemic regions report lower COVID-19 morbidity and mortality. Here, we show that lung remodeling from a prior infection with a lung migrating helminth, *Nippostrongylus brasiliensis*, enhances viral clearance and survival of human-ACE2 transgenic mice challenged with SARS-CoV-2 (SCV2). This protection is associated with a lymphocytic infiltrate including an increased accumulation of pulmonary SCV2-specific CD8+ T cells and anti-CD8 antibody depletion abrogated the *N. brasiliensis*-mediated reduction in viral loads. Pulmonary macrophages with a type-2 transcriptional signature persist in the lungs of *N. brasiliensis* exposed mice after clearance of the parasite and establish a primed environment for increased antigen presentation. Accordingly, depletion of macrophages ablated the augmented viral clearance and accumulation of CD8+ T cells driven by prior *N. brasiliensis* infection. Together, these findings support the concept that lung migrating helminths can limit disease severity during SCV2 infection through macrophage-dependent enhancement of anti-viral CD8+ T cell responses.

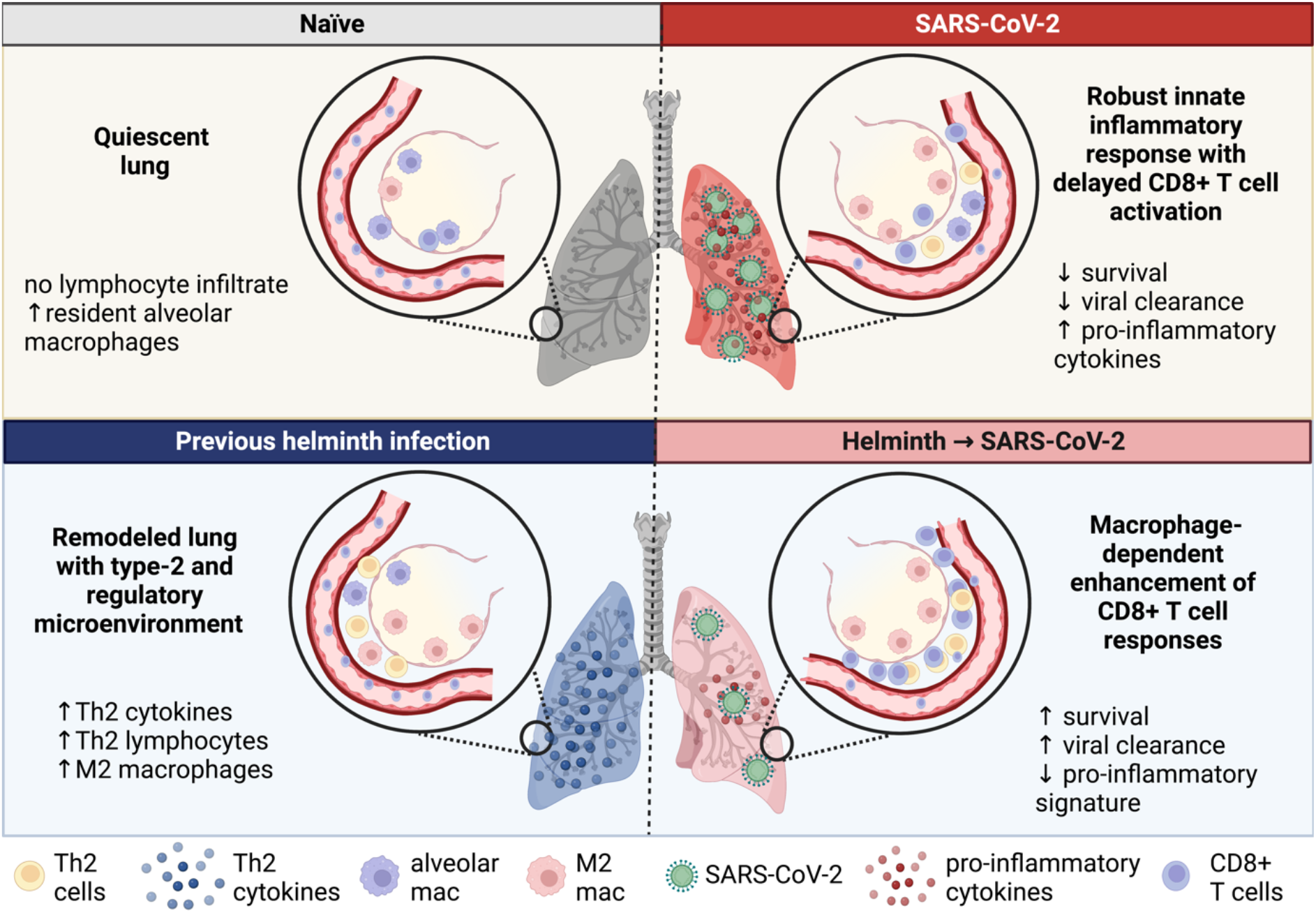

## Introduction

Regions with a high prevalence of helminth infections report lower morbidity and mortality related to COVID-19 (1, 2), raising the possibility that worm infections can modulate SARS-CoV-2 (SCV2) disease outcomes. Indeed, a small hospital-based study in Ethiopia reported that COVID-19 patients co-infected with helminth parasites have a lower risk of developing severe disease (3). Nevertheless, there are conflicting views on the impact of helminth infections on COVID-19 (4–9), and the effects of worm infection on the pulmonary response to SCV2 remains unclear.

Previous studies have shown that helminth infections can both worsen (10, 11) or be beneficial for the outcome of other viral infections (11–15). The immune response, tissue tropism and the timing of viral exposure in relation to the helminth life cycle (11, 16) are all factors potentially affecting viral disease outcomes. Notably, prior but not concurrent infection with a lung traversing helminth is associated with beneficial outcomes following influenza challenge in mice(16) and adults from helminth endemic countries would likely have had prior exposure to worms as children (17–19).

Helminth infection typically induces an innate and adaptive type-2 and regulatory response (20–22) as well as cellular phenotypes such as alternatively activated M2 macrophages (23). Importantly, these Type-2 and regulatory profiles can persist even after worm clearance (24–29) and thus may influence tissue responses to subsequent immunological challenges. Furthermore, helminth infections can also induce expansion of virtual memory CD8+ T cells, which can promote viral clearance (12, 30, 31).

Here, we investigated the impact of lung remodeling from prior helminth infection on SCV2 disease pathogenesis using the K18-hACE2 mouse model (32, 33). We show that prior infection by the lung migrating helminth, *Nippostrongylus brasiliensis*, improves disease outcomes following SCV2 challenge and enhances viral clearance. Both CD8+ T cells and pulmonary macrophages were found to be required for *N. brasiliensis*-conferred protection against SCV2, revealing a pathway for long-lived helminth-elicited responses in promoting pulmonary host resistance to subsequent viral challenge.

## Results

### Previous infection with *N. brasiliensis* enhances viral clearance and protects K18-hACE2 mice against SCV2 driven lethality

*N. brasiliensis* migrating larvae cause extensive lung tissue damage that is rapidly repaired (34, 35). The parasite is then cleared in wildtype mice 7-10 days post infection. To investigate if lung remodeling impacts SCV2 disease, SCV2-susceptible K18-hACE2 mice were infected with 500 *N. brasiliensis* larvae subcutaneously (s.c.) and rested to naturally clear and recover from their worm infections for a 28-day period. The animals were then challenged intranasally (i.n.) with a lethal dose of SCV2 WA/2020 and monitored for weight change and survival (**Fig1A**). Similar to *N. brasiliensis*-naïve control mice, animals that had previously been infected with *N. brasiliensis* rapidly lost weight following SCV2 challenge (**Fig1B**). However, prior *N. brasiliensis* exposure conferred a significant survival benefit, with 60% of *N. brasiliensis* exposed mice surviving SCV2 infection compared to just 20% of controls (**Fig1C**).

**Figure 1:**
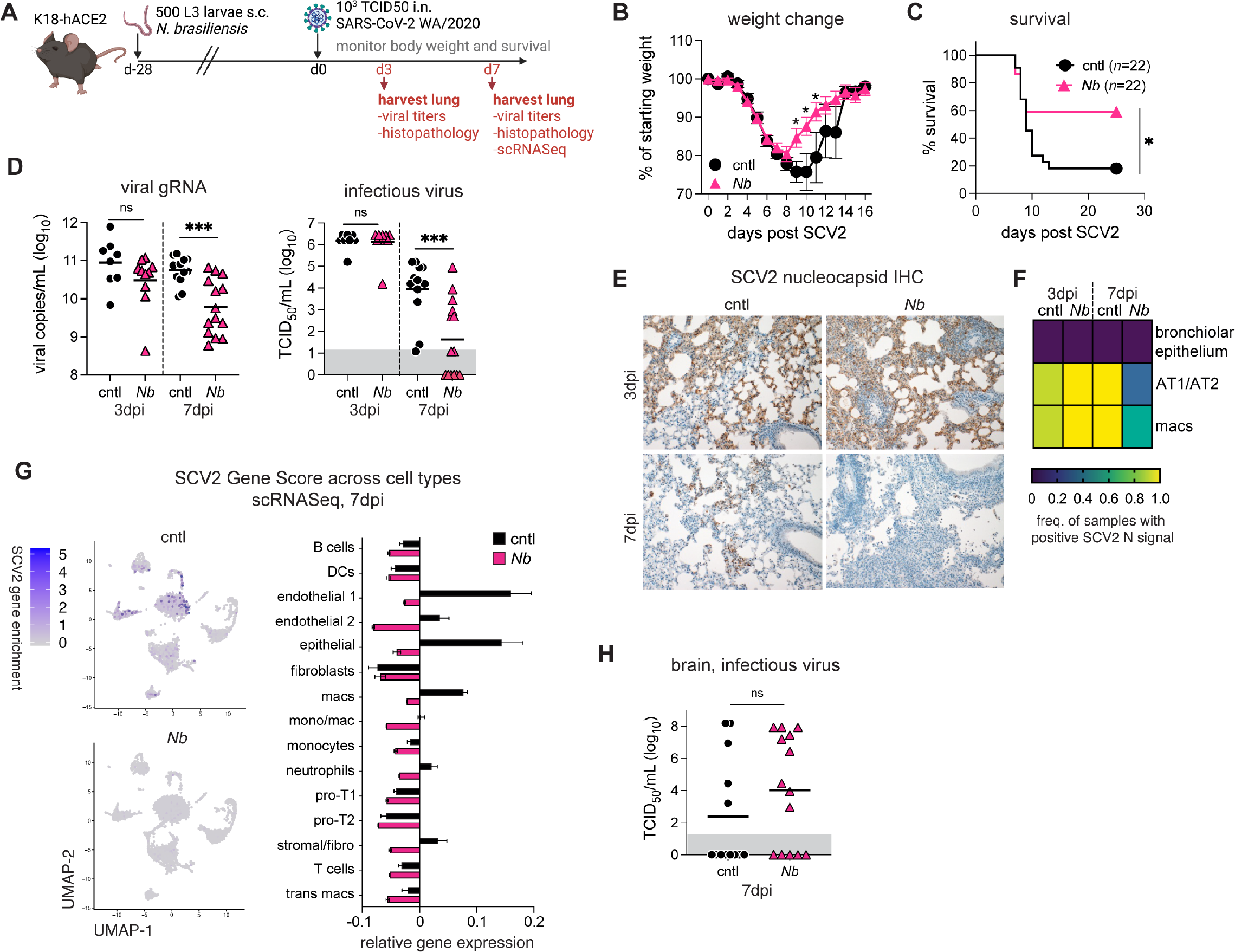
Previous infection with *N. brasiliensis* enhances viral clearance and protects K18-hACE2 mice against SCV2 driven lethality. K18-hACE2 mice were infected with 500 L3 *N. brasiliensis* (*Nb*) larvae subcutaneously (s.c.) or left uninfected (cntl). After 28 days, all animals were challenged intranasally (i.n.) with 10^3^ TCID50 SCV2. (**A**) Schematic of experimental protocol. (**B**) Weight change following SCV2 infection, shown as percentage of average weight on d-1 and d0. *n*=22 mice/group; 3 independent experiments. Statistical significance was determined by unpaired *t*-test. Mean ± SEM. (**C**) Kaplan-Meier curve of animal survival following SCV2 challenge. *n*=22 mice/group; 3 independent experiments. Statistical significance was assessed by Mantel-Cox test. (**D**) Lungs were collected 3 or 7d after SCV2 challenge and assessed for viral load by qPCR or TCID50 assay. *n*=8-14 mice/group; 2-3 independent experiments. Statistical significance was determined by Kruskal-Wallis test with Dunn’s post-test. Geometric mean is shown. Gray box indicates values below limit of detection. (**E**) Representative images of lung tissue sections probed with anti-SCV2 nucleoprotein antibody at 3 or 7d post SCV2 challenge. (**F**) Heat map showing the frequency of animals with detectable SCV2 N protein in bronchiolar epithelial cells, pneumocytes (AT1/AT2) or macrophages as determined by a board-certified veterinary pathologist. *n*=7-10 mice/group; 2 independent experiments. (**G**) Feature plot showing scRNA-seq viral transcripts made up of “orf1ab", “S", “orf3a", “E", “M", “orf6", “orf7a", “orf8", “N", “orf10” viral genes and bar plot showing the gene expression of the SCV2 viral genes in the different cell types. *n*=pool of 3-4 mice/group. (**H**) Brains were collected 3 or 7d after SCV2 challenge and assessed for viral load by TCID50 assay. *n*=8-14 mice/group; 2-3 independent experiments. Statistical significance was determined by Kruskal-Wallis test with Dunn’s post-test. Geometric mean is shown. Gray box indicates values below limit of detection. ns *p*>0.05; * *p*<0.05; *** *p*<0.001

We assessed viral loads in the lungs of *N. brasiliensis* and control animals 3 or 7 days after SCV2 challenge to reflect the peak viral load (3dpi) and when disease outcomes bifurcate between control and *N. brasiliensis* animals (7dpi). Viral loads in lung homogenates were similar between groups at 3dpi by qPCR measurement of SCV2 genomic E expression as well as tissue culture infectious dose-50 (TCID50) assays (**Fig1D**). However, *N. brasiliensis* mice showed significantly lower viral loads at 7dpi when compared to controls (**Fig1D**). Immunohistochemical analysis of lung sections showed that SCV2 nucleocapsid immunoreactive pneumocytes (AT1/AT2) and macrophages were less common in lung tissue sections at 7dpi, whereas no difference was observed at 3dpi (**Fig1E-F**). We performed single-cell RNAseq of lung cells at 7dpi and mapped SCV2 transcripts as a gene enrichment score incorporating expression of *orf1ab*, *S*, *orf3a*, *E*, *M*, *orf6*, *orf7a*, *orf8*, *N*, *orf10* viral genes, and found that SCV2 genes were less abundant in endothelial cells, epithelial cells, neutrophils, stromal cells and macrophages from *N. brasiliensis* mice (**Fig1G**). Viral loads in the brain were similar (**Fig1H**), arguing against differential central nervous system involvement, which can contribute to mortality (36–39). Together, these data suggested that prior *N. brasiliensis* infection enhances the clearance of SCV2 in the lung to promote host survival.

### CD8+ T cells are necessary for enhanced viral clearance in *N. brasiliensis* remodeled lungs

The reduced SCV2 titers at 7dpi but not at 3dpi suggested improved adaptive immune responses, rather than enhanced innate responses against viral establishment. Indeed, histopathology showed that lymphocytic inflammation in lung parenchyma of mice previously infected with *N. brasiliensis* was more pronounced than control animals (**FigS2A-B**). Flow cytometric analysis revealed that the percentage and number of lung CD8+ T cells is markedly higher at 7dpi in *N. brasiliensis* exposed mice as compared to controls (**Fig2A**). While total CD8+ T cell numbers were similar between groups at 3dpi (**Fig2A**), a higher proportion of these CD8+ T cells are localized within the lung tissues in *N. brasiliensis* mice (**Fig2B**), as assessed by intravenous (i.v.) staining of CD45+ cells, used to discriminate cells in the pulmonary vasculature from those in the lung interstitium or airways (40). When we stained for SCV2 spike (S)-specific cells using a S_539-546_ tetramer, we found that S-specific T cells were absent at 3dpi and increased in frequency by 7dpi (**Fig2C**), which could be important for protection (41, 42). These results suggest that *N. brasiliensis* exposure increases recruitment and/or accumulation of SCV2 specific CD8+ T cells in the lung tissue following SCV2 infection.

**Figure 2:**
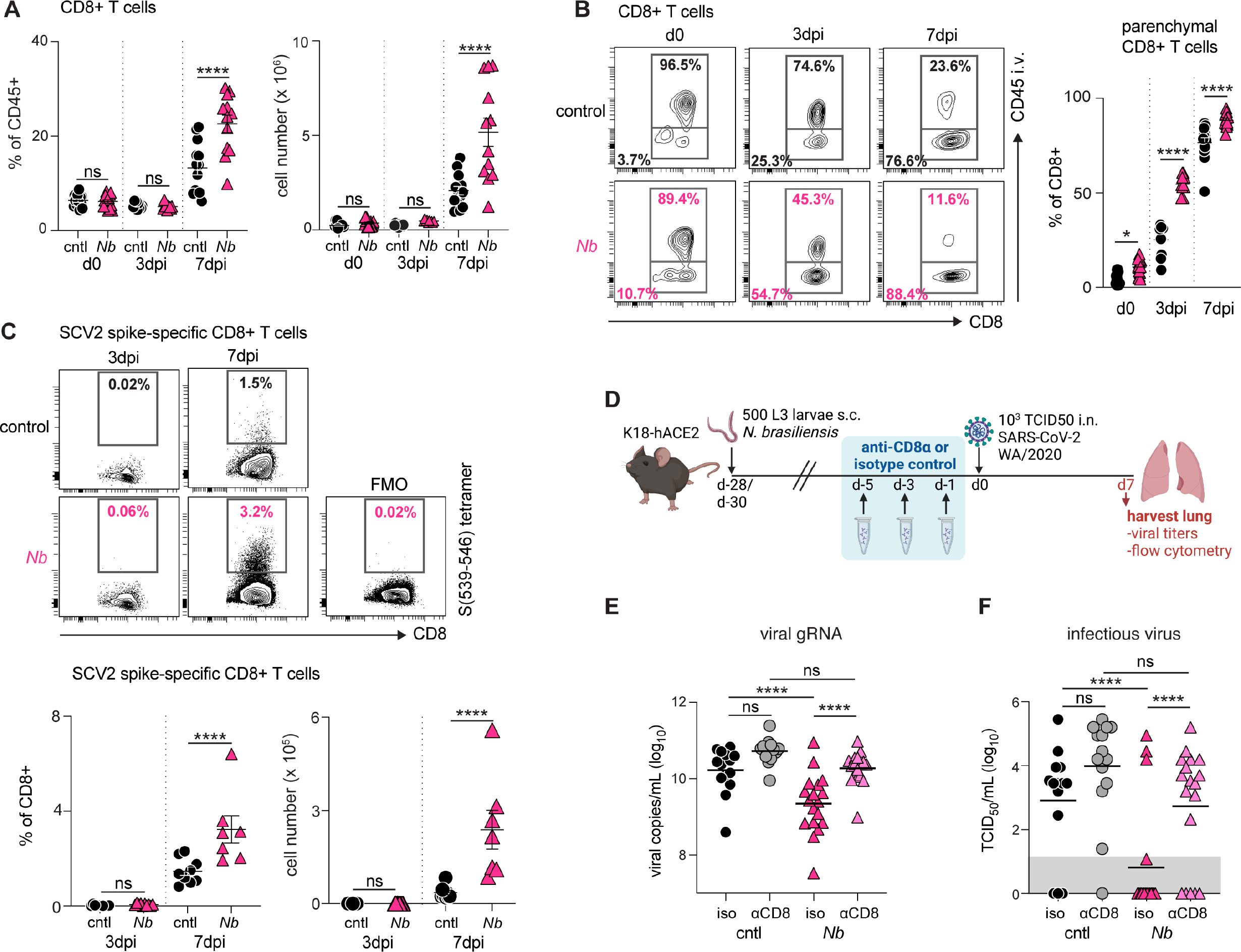
Previous *N. brasiliensis* infection amplifies CD8+ T cell responses following SCV2 challenge and depletion of CD8+ T cells abrogates *N. brasiliensis-* mediated control of viral loads. (**A-D**) K18-hACE2 mice were infected with 500 L3 *N. brasiliensis* (*Nb*) larvae s.c. or left uninfected (cntl). After 28 days, animals were challenged i.n. with 10^3^ TCID50 SCV2. Control animals did not receive SCV2 challenge (d0). At 3 or 7d post SCV2, lungs were harvested and processed for flow cytometric analysis. *n*=7-15 mice/group; 2-3 independent experiments. Statistical significance was assessed using a linear mixed-effects model with pairwise comparison using JMP software. Data are displayed as mean ± SEM. (**A**) Frequency and number of total CD8+ T cells. (**B**) Frequency of parenchymal (CD45 i.v.-) vs. vascular (CD45 i.v.+) CD8+ T cells. (**C**) Frequency and number of CD8^+^ T cells positive for the SCV2-specific S_(539-546)_ tetramer. (**D-F**) K18-hACE2 mice were inoculated with 500 *N. brasiliensis* larvae by s.c. injection at d-28. Mice were then treated with either anti-CD8α or rat IgG2b isotype control on d-5, d-3, d-1 prior to SCV2 challenge on d0. Lung tissue was harvested at 7dpi. *n*=11-17 mice/group; 3 independent experiments. Statistical significance was assessed using a linear mixed-effects model with pairwise comparison using JMP software. Data are displayed as mean ± SEM. (**D**) Schematic of experimental protocol. Viral loads as measured by qPCR (**E**) or TCID50 assay (**F**). Geometric mean is shown. Gray box indicates values below limit of detection. ns *p*>0.05; * *p*<0.05; **** *p*<0.0001

We next treated animals with an anti-CD8α depleting antibody or an isotype-matched control on days −5, −3 and −1 leading up to intranasal SCV2 challenge (d0) (**Fig2D**). The antibody treatment successfully depleted CD8+ T cells in the lungs during SCV2 infection (**FigS2C**) and led to a significant increase in viral burden in *N. brasiliensis* mice, with viral gRNA and infectious particles being restored to levels observed in control animals (**Fig2E-F**). Lung viral loads tended to be higher with anti-CD8α treatment in control mice at 7dpi, but this did not reach statistical significance. Hence, lung remodeling by prior *N. brasiliensis* infection confers anti-SCV2 protection in a CD8+ T cell dependent manner.

### *N. brasiliensis* infection results in long-term alterations in pulmonary CD4+ T cells and macrophages

To understand how lung remodeling by *N. brasiliensis* shapes the immune environment prior to exposure to SCV2, we performed single cell RNA sequencing (scRNA-seq) analysis on lung cells at 28 days post infection with *N. brasiliensis* (i.e. the time of SCV2 challenge). Seurat clustering (43) revealed 20 cell clusters (**Fig3A**) which were identified by singleR (44) and marker genes (**FigS3A and Table S1**). This analysis showed a notable enrichment in lymphocytes, dendritic cells and alternatively activated macrophages in *N. brasiliensis* exposed lungs (**Fig3B**). Higher expression of Type 2 cytokines (IL-4, IL-5 and IL-13) was detectable transcriptionally based on scRNA-seq (**Fig3C**) and as proteins by a multiplex assay (**Fig3D**). *Il4* was primarily detected in granulocytes, *Il5* in lymphocytes and *Il13* transcripts were found in both subsets (**FigS3B**). Spectral cytometry revealed an increase in the number and frequency of eosinophils (**FigS3C**) and Group 2 innate lymphoid cells (ILC2s) (**FigS3D**) in *N. brasiliensis* exposed mice, hence eosinophils are the likely source of *Il4* and ILC2s the source of *Il5* and *Il13* transcripts (45–47). The multiplex cytokine analysis also revealed that IL-1β, IL-18, TNFα, CXCL10, GM-CSF and IL-12p70 levels were all significantly higher in *N. brasiliensis* mice (**Fig3D**). *N. brasiliensis* exposed lungs also had higher frequencies and numbers of CD4^+^ T cells (**Fig3E-F**), which are activated (CD44^hi^, **Fig3E and FigS3E**) and resident within the tissue parenchyma (CD45 i.v.-) (**Fig3G**). More CD4+ cells are polarized towards a Th2 (GATA3^+^) or Treg (FoxP3^+^) phenotype (**Fig3E, Fig3G and FigS3F**) and produce IL-10 following stimulation with PMA/ION after *N. brasiliensis* exposure (**FigS3G**). By scRNA-seq, the CD4+ lymphocyte cluster expresses more transcripts of Th2 markers including *Il1rl1, Gata3, Icos, Malt1, Maf, Rbpj* and *Areg* (**Fig3E**) and may also be another source of the Type 2 cytokine transcripts observed in the scRNA-seq data (48). These results suggest that following remodeling by *N. brasiliensis* infection, the lung retained a primed inflammatory and a long-lasting Th2 signature.

**Figure 3:**
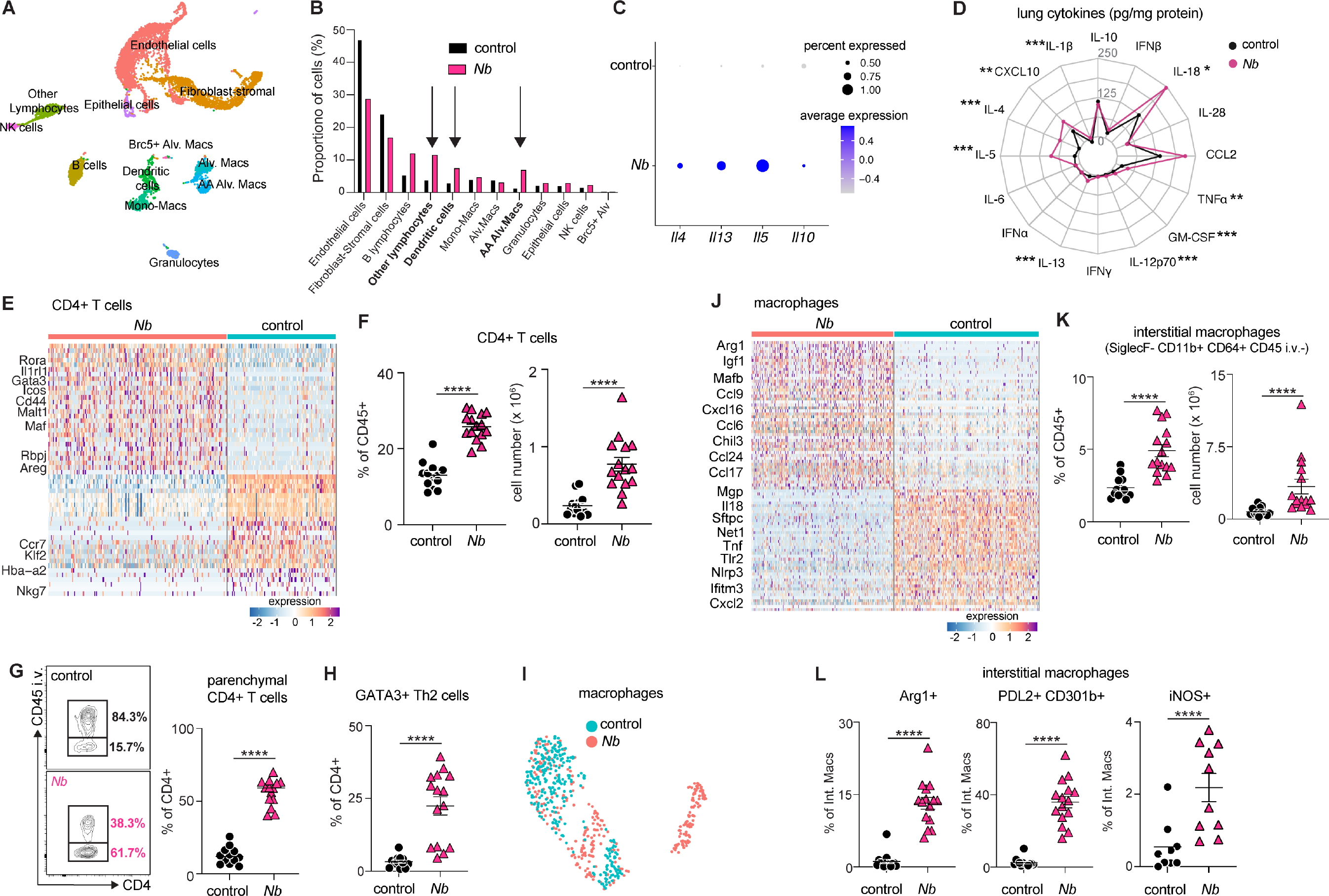
Previous *N. brasiliensis* infection results in long-term alterations in the pulmonary CD4+ T cell and macrophages compartments. K18-hACE2 mice were infected with 500 L3 *N. brasiliensis* (*Nb*) larvae s.c. or left uninfected (control). After 28 days, lungs were harvested and processed for scRNAseq (*n*=pool of 4-5 mice/group), flow cytometric analysis or multiplex cytokine assay (*n*=14-15 mice/group; 3 independent experiments). (**A**) UMAP visualization of scRNASeq data of control (*n* = 5385 cells) and *Nb* (*n* = 2897 cells) identifying 20 distinct cell clusters. (**B**) Frequency of cell types identified in (A) separated by treatment group. (**C**) Normalized expression of *Il4*, *Il5*, *Il13* and *Il10* transcripts across all cell types. (**D**) Radar plot showing mean protein levels (pg/mg total protein) of cytokines and chemokines in different groups measured by multiplex assay in whole lung homogenate. Statistical significance was determined by unpaired Student’s *t*-test between the two groups for each cytokine/chemokine assessed. (**E**) Differential expression analysis of the CD4+ T cell cluster showing the key genes out of the top 50 DEGs between *Nb* and control groups. Each column represents an individual cell. (**F-H**) Flow cytometric determination of the (**F**) frequency and number of total CD4^+^ T cells, (**G**) frequency of parenchymal/airway (CD45 i.v.-) vs. vascular (CD45 i.v.+) CD4^+^ T cells and (**H**) frequency of GATA3^+^ CD4^+^ Th2 cells. (**I**) UMAP of re-clustered alveolar macrophages subset visualized by treatment group. (**J**) Differential expression of alveolar macrophages showing the key genes out of the top 50 DEGs between *Nb* and control. Each column represents an individual cell. (**K-L**) Flow cytometric determination of the (**K**) frequency and number of Siglec F^−^ CD11b^+^CD64^+^ CD45 i.v.^−^ interstitial macrophages and (**L**) their expression of Arg1, PDL2, CD301b and iNOS. Statistical significance was assessed using a linear mixed-effects model with pairwise comparison using JMP software. ns *p*>0.05; * *p*<0.05; ** *p*<0.01; *** *p*<0.001; **** *p*<0.0001

In addition to Th2 cells, antigen presenting cells (APCs) are also expanded and skewed towards a Type 2 phenotype 28 days after *N. brasiliensis* infection (**Fig3B and FigS3H)**. In particular, macrophages with an alternative activation phenotype expressing arginase 1 (*Arg1*), mannose receptor 1 (*Mrc1*), chitinase-like protein 3 (*Chil3*) and matrix metalloproteinase (*Mmp12*)(49–51) **(Table S1)** are more abundant in *N. brasiliensis* exposed lungs (**Fig3B**). Re-clustering of the alveolar macrophage populations in the scRNA-seq dataset revealed striking differences (**Fig3I and FigS3I**), with many top 50 differentially expressed genes associated with tissue remodeling and alternative activation (*Arg1, Ctsk, Ctss, Igf1* and *Chil3*), as well as cellular chemotaxis and migration (*Ccl9, Cxcl16, Ccl24, Trem2* and *Ccl17)* (**Fig3J, FigS3J and Table S2**). Consistent with these data, spectral cytometry showed that more of the CD45^+^SiglecF^+^CD11c^+^ alveolar macrophages from *N. brasiliensis* exposed mice expressed arginase 1 (Arg1), PDL2 and CD301b (52–54) (**FigS3K**), although the numbers were similar between groups (**FigS3L)**. Furthermore, we observed more tissue resident CD11b^hi^ interstitial macrophages (**Fig3K**) (55), which also express more alternatively activated macrophage markers, Arg1, PDL2 and CD301b (**Fig3L**). Together, these results suggest that *N. brasiliensis* infection leads to a long-lasting phenotypic and transcriptional changes, with pulmonary immune cells skewed towards a Type-2 or alternative activation phenotype that persists even after worm clearance.

### Pulmonary macrophages are required for enhanced viral clearance and CD8+ T cell responses

We next performed scRNASeq on lung cells 7 days after SCV2 infection in *N. brasiliensis* exposed and naïve control mice and identified 15 cell clusters, including 3 macrophage clusters (**Fig4A, FigS4A-C and Table S3**). As previously noted in acute inflammatory responses (56), we observed a loss of alveolar macrophages at day 7 post SCV2 challenge, with cells expressing characteristic alveolar macrophage genes making up a smaller percentage of the monocyte-macrophage compartment as opposed to what is seen in before SCV2 infection (**Fig4A and FigS4D**). After sub-setting and re-clustering of the monocyte/macrophage compartment, two major macrophage populations can be separated that are differentially enriched in *N. brasiliensis* exposed and control lungs, as defined by expression of *Sec61a1* (*N. brasiliensis*) and *Scgb1a1* (control) (**Fig4B and 4C**). While transcriptionally distinct, both these populations expressed some genes associated with an alveolar macrophage phenotype, suggesting that they may be inflammatory macrophages transitioning into alveolar macrophages (**FigS4D**). By mapping SCV2 transcripts, *Scgb1a1* lung macrophages from control mice had higher levels of viral RNA (**Fig4D**), consistent with the higher virus loads in the lungs of these mice (**Fig1**). Genes involved in antigen processing and presentation were found to be enriched in the *Sec61a1* macrophage populations from *N. brasiliensis* exposed mice (**Fig4E**). In contrast, the *Scgb1a1* macrophages expressed pro-inflammatory cytokine response genes, such as *Ccl2, Saa3, Ccl7*, and *Cxcl13*, which may be driven by the persisting viral load in these mice.

**Figure 4:**
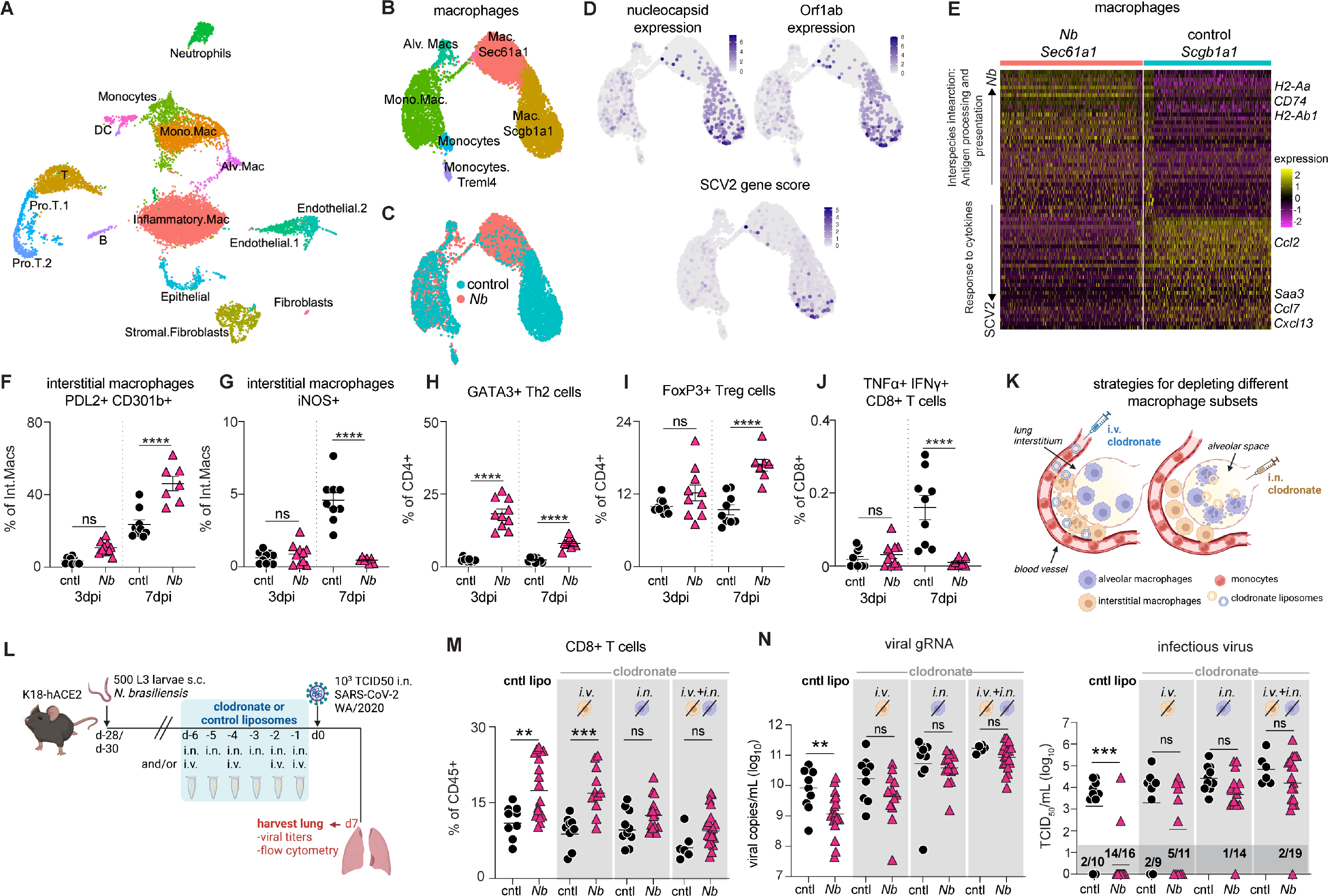
Depletion of pulmonary macrophages abrogates *N. brasiliensis* enhanced viral clearance while diminishing CD8+ T cells responses. (**A-J**) K18-hACE2 mice were infected with 500 L3 *N. brasiliensis* (*Nb*) larvae s.c. or left uninfected (cntl). After 28 days, animals were challenged i.n. with 10^3^ TCID50 SCV2. At 7d post SCV2, lungs were harvested and processed for scRNASeq (*n*=pool of 3-4 mice/group) or flow cytometry (*n*=7-15 mice/group; 2-3 independent experiments). UMAP visualization of (**A**) of control (*n* =6402 cells) and *Nb* (*n*=6930 cells), (**B**) re-clustered monocytes-macrophages cluster, (**C**) overlay of the monocyte-macrophage cluster from the different experimental groups and (**D**) macrophages with overlay of viral gene expression (N and Orf1ab) and SCV2 gene score encompassing all viral genes. (**E**) Differential expression analysis of *Sec61a1* (*Nb*) and *Scgb1a1* (control) macrophages showing the top 50 DEGs. Enriched pathways are listed on the lefthand side. Each column represents an individual cell. (**F-J**) Flow cytometric determination of the frequency of (**F**) PDL2^+^ CD301b^+^ M2 macrophages, (**G**) iNOS^+^ M1 macrophages, (**H**) GATA3^+^ CD4^+^ Th2 cells, (**I**) FoxP3^+^ CD4^+^ Tregs and (**J**) TNFα^+^ IFNγ^+^ CD8+ T cells. Statistical significance was assessed using a linear mixed-effects model with pairwise comparison using JMP software. Data are displayed as mean ± SEM. (**K-N**) K18-hACE2 mice were inoculated with 500 *Nb* larvae by s.c. injection at d-28. Mice were then treated with either clodronate liposomes or control liposomes by i.n. and/or i.v. administration from d-6 to d-1 prior to SCV2 challenge on d0. Lungs were harvested at 7dpi. *n*=6-19 mice/group; 2 independent experiments. Statistical significance was determined by Mann-Whitney test. (**K**) Illustration of strategy for targeting interstitial and airway macrophages for depletion. (**L**) Schematic of experimental protocol. (**M**) Frequency of lung CD8+ T cells as determined by flow cytometry. Data are mean ± SEM. (**N**) Viral loads measured by qPCR and TCID50 assay. Geometric mean is shown. Dark gray box indicates values below limit of detection. ns *p*>0.05; * *p*<0.05; ** *p*<0.01; *** *p*<0.001; **** *p*<0.0001

Consistent with the scRNAseq data, spectral cytometry showed that even 7 days after an acute viral infection, macrophages from the *N. brasiliensis* exposed mice retain a Type-2 phenotype with higher expression of alternatively activation markers (PDL2 and CD301b) (**Fig4F**) and decreased expression of iNOS (**Fig4G**) and a higher proportion of Th2 (GATA3^+^) (**Fig4H**) and Treg (FoxP3^+^) CD4+ cells (**Fig4I**). CD8+ T cells in the lungs of *N. brasiliensis* exposed mice produce less pro-inflammatory cytokines such as IFNγ and TNFα (**Fig4J, FigS4E**), and by scRNA-seq showed lower expression of genes associated with cytotoxic activity (e.g., *Gzmb, Gzma*) and inflammation (*Cxcl10*, *Cxcl9*, *Tnfrsf9*) (**FigS4F**). IFNγ and TNFα protein levels measured in lung homogenates from the same animals are also reduced (**FigS4G and S4H**). Notably, S-specific CD8+ cells also produce less Granzyme B (**FigS4I**) in lungs of *N. brasiliensis* exposed mice. The reduced CD8+ T cell cytokine response in the *N. brasiliensis* exposed mice could be due to the lower viral loads or may reflect an increased regulatory response by macrophages and CD4+ cells. Together, these data show that at 7dpi with SCV2, macrophages from *N. brasiliensis* exposed mice exhibit up-regulated antigen processing and presentation genes that may contribute to the enhanced CD8+ T cell response required for effective viral clearance, while there is a reduced pro-inflammatory module associated with COVID-19 disease severity (57).

We next selectively depleted the interstitial and/or alveolar macrophage subsets *in vivo* with clodronate liposomes administered via different routes (i.v. to remove interstitial and/or i.n. to remove alveolar cells) over the week preceding SCV2 challenge (58, 59) (**Fig4K-L**). We stopped clodronate treatment 1 day before viral challenge because circulating monocytes have been implicated in SCV2 control (60). *N. brasiliensis* exposed mice depleted of alveolar macrophages alone via the i.n. route no longer had significantly more CD8+ T cell responses, which in contrast remained elevated in *N. brasiliensis* exposed mice treated with the control liposomes or depleted of interstitial macrophages by i.v. treatment (**Fig4M and FigS4J**). Hence, macrophages in the alveoli are required for the enhanced CD8+ response against SCV2 in *N. brasiliensis* exposed mice, whereas interstitial macrophages are dispensable. The combination of i.v. and i.n. clodronate administration also reversed the enhanced CD8+ T cell response (**Fig4M**). When we examined viral loads by RT-qPCR and TCID50 assays (**Fig4N, FigS4K and S4L**), control liposome treated mice had significantly reduced viral loads after *N. brasiliensis* exposure as expected, however all the clodronate liposome treated animals no longer exhibited differences in viral load (**Fig4N, FigS4K and S4L**). Hence, while alveolar macrophages may be more important for CD8+ T cell responses, interstitial macrophages may also be required for controlling viral loads. However, it should be noted that 5 out of 11 mice in the i.v. treated group had undetectable viral loads by the TCID50 assay, whereas only 1 of 14 in the i.n. treated group and 2 of 19 in i.v.+i.n. treated group had undetectable viral loads (**Fig4N, FigS4K and S4L**). Hence the mice depleted of interstitial macrophages have more of an intermediate phenotype. Together, these results demonstrate that helminth-primed pulmonary macrophages are critical for enhancing CD8+ T cell responses and viral clearance following subsequent SCV2 infection and that macrophages located in the alveoli may be more important for this function than interstitial macrophages.

## Discussion

We observed that previous infection of mice with the lung migrating helminth parasite, *N. brasiliensis*, accelerates viral clearance and reduces mortality from subsequent SCV2 infection. Mechanistically, we find that this protection is mediated by enhanced recruitment and/or expansion of virus specific CD8+ T cells and that pulmonary macrophages altered by helminth infection are required for the augmented CD8+ T cell response and viral clearance. While these observations were made in a mouse model, our results are consistent with a hospital study in Ethiopia whereby patients co-infected with intestinal parasites, including helminths, had lower odds of developing severe COVID-19 (3). Nevertheless, there are still no epidemiologic data addressing whether helminth infection increases susceptibility to SARS-CoV-2 infection itself as opposed to COVID-19 disease progression.

The role of Type-2 cytokines in viral immunity is complex. Type-2 responses have been associated with worse COVID-19 disease outcomes in mice and humans. Patients with severe COVID-19 have increased levels of IL-5, IL-13, IgE and eosinophils in circulation (61) and monoclonal antibody blockade of IL-4 and IL-13 receptor signaling improved their prognosis (62). IL-4 can inhibit anti-viral immunity against influenza (63) and respiratory syncytial virus (RSV) (64). Nevertheless, helminth infection in mice (with the intestinal parasite *H. polygyrus*) protects against pulmonary inflammation from RSV infection via type 1 interferon enhanced viral clearance (15). In contrast, the same parasite increases susceptibility to flavivirus infection, increasing both viral load and disease mortality (65), and can reactivate latent gamma herpesvirus through IL-4 driven STAT6 signaling (66). *N. brasiliensis*, the helminth that we show here protects against SARS-CoV-2 infection, can also exacerbate viral-induced epithelial ulceration and pathology in herpes simplex virus (HSV)-2 infection (67). However, there is also strong evidence that IL-4 driven by helminth infections can enhance CD8+ T cell responses (68) and promote the rapid generation of antigen-specific T responses during viral infection (12). While our data demonstrate that a pulmonary environment primed by larval migration to adopt a Type 2 microenvironment enhances viral specific CD8+ T cells and viral clearance, we have not directly shown that Type 2 cytokines are required for the enhanced protection.

Our results demonstrate that helminth-primed macrophages are important for the generation of the augmented CD8+ T cell response as well as accelerated viral clearance seen in helminth infected mice. While the precise immunologic mechanism governing this outcome is unclear, lung remodeling following larval migration clearly promotes an activated macrophage profile that is still present 28 days after *N. brasiliensis* infection. Monocytes are recruited to the lung after *N. brasiliensis* larval migration and differentiate into an alveolar macrophage phenotype (69). These monocyte-derived alveolar macrophages express type 2-associated markers and can mediate enhanced helminth killing in a secondary infection (69). The findings presented here indicate that these monocyte-derived alveolar macrophages also mediate enhanced anti-viral clearance through modulating protective T cell responses. These pulmonary macrophages are characterized by the expression of pro-inflammatory cytokines and chemokines such as *Cxcl16* and *Ccl17*, which are important for the rapid recruitment of T cells into the lung parenchyma (70–72). In addition, the increased antigen presentation profile seen in the helminth primed macrophages following exposure to SCV2 is likely important for promoting increased activation and stimulation of the CD8+ T effector cells required for effective viral clearance (73). Reciprocal interactions between antigen-presenting cells and CD4+ T cells have been shown to play a role in boosting anti-viral CD8+ T cell responses (74). While we did not directly address this possibility in the present study, we do show that the helminth macrophage antigen presentation profile consists of molecules associated with MHCII presentation and that prior *N. brasiliensis* infection greatly enriches CD4+ T cells in the lung. Therefore, such processes may be involved in driving the augmented anti-SCV2 CD8+ T cell response observed in mice that have recovered from a helminth infection. These scenarios are consistent with data from COVID-19 patients indicating that antigen presentation modules are enriched in macrophage populations from individuals with favorable disease outcomes (75).

In addition to mediating enhanced recruitment and activation of CD8+ cells, helminth-primed macrophages with an alternatively activated phenotype are known to be critical for tissue repair following inflammation (76). Tissue remodeling factors such as *Arg1, Ctsk, Ctss, Igf1* and *Chil3* are up-regulated in macrophages from *N. brasiliensis*-infected mice and are associated with resolution of inflammation (76–81). Activation of these macrophages can be amplified following re-infection (76) and may promote recovery and repair of the lung tissue following exposure to a highly inflammatory insult like SCV2. Together with the increased frequency of regulatory T cells and the accelerated clearance of virus in helminth exposed mice, alternatively activated macrophages may contribute to the dampening of pro-inflammatory modules observed later in disease thereby promoting recovery and less severe outcomes in mice previously infected with *N. brasiliensis*.

Increased protection against SCV2 in an immunologically primed pulmonary environment is not a unique feature of Type 2 immunity and helminth infection. Prior or concurrent bacterial infection of the lung can also alter the response to SCV2 and improve disease outcomes in mice (82–84) (Paul Baker and Katrin Mayer-Barber, personal communication). Indeed, the increased accumulation of both innate and adaptive immune cells in lung tissue after inflammatory events may have generalized protective effects against SCV2 challenge. The data presented here implicate macrophages as an essential component of this primed non-specific protection. Understanding these different tissue priming events can help to uncover mechanisms of host resistance to SCV2 and at the population level help understand how prior infection with unrelated pathogens could influence COVID-19 in different endemic settings. At a more general level, the findings presented here provide a striking example of how an individual’s immunological history can modulate their subsequent response to unrelated pathogen exposure.

## Methods

### Mice

B6.Cg-Tg(K18-ACE2)2Prlmn/J hemizygous (JAX34860) mice were purchased from The Jackson Laboratory (Bar Harbor, ME) and were housed under specific pathogen–free conditions with *ad libitum* access to food and water. Animals were randomly assigned to sex- and age-matched experimental groups. All studies were conducted in AALAC–accredited Biosafety Level 2 and 3 facilities at the NIAID, National Institutes of Health (NIH) in accordance with protocols approved by the NIAID Animal Care and Use Committee.

### Virology

SARS-CoV-2 strain USA-WA1/2020 (BEI Resources) was propagated in Vero-TMPRSS2 cells (kindly provided by Dr. Jonathan Yewdell, NIAID) under BSL3 conditions in DMEM medium supplemented with Glutamax and 2% FCS. At 48h post inoculation, culture supernatant and cells were collected and clarified by centrifugation for 10 min at 4°C. Supernatant was collected, aliquoted and frozen at −80°C. Viral titers were determined by TCID_50_ assay in Vero E6 cells (ATCC CRL-1586) using the Reed and Muench calculation method. Full genome sequencing was performed at the NIAID Genomic Core (Hamilton, MT). The virus stock used in this study contained 2 single-nucleotide polymorphisms from the reference sequence MN985325.1: T7I (M), S194T (N).

### Infections and treatments

Mouse-adapted *N. brasiliensis* was maintained by serial passage intermittently through C57BL/6 mice and STAT6-KO mice, as described previously (85). Animals were infected with 500 third stage *N. brasilienesis* larvae by subcutaneous injection.

For CD8+ T cell depletion studies, 200μg anti-CD8α (YTS 169.4) or rat IgG2b isotype control (LTF-2) was administered by intraperitoneal injection on days −5, −3 and −1 as indicated in the text and figures. Antibodies were stored at 4°C until use and diluted in InVivoPure dilution buffer just prior to administration. Antibodies and buffers were from BioXCell.

For macrophage depletion studies, liposome-encapsulated clodronate or empty control liposomes (both from Encapsula NanoSciences) were administered by intranasal instillation (175μg/dose) and/or intravenous injection (500μg/dose). Intranasal administrations were performed daily from d-6 to d-1. Intravenous treatments were delivered on days −6, −4, −2 and −1 (59). Clodronate and control liposomes were stored at 4°C and administered undiluted to animals at the indicated time points.

SCV2 infections were performed under BSL3 containment. Animals were anesthetized by isoflurane inhalation and a dose of 10^3^ TCID_50_/mouse SCV2 WA/2020 was administered by intranasal instillation. Following infection, mice were monitored daily for weight change and clinical signs of disease by a blinded observer who assigned each animal a disease score based on the following criteria: 0) no observable signs of disease; 1) hunched posture, ruffled fur and/or pale mucous membranes; 2) hunched posture and ruffled fur with lethargy but responsive to stimulation, rapid/shallow breathing, dehydration; 3) moribund.

### Determination of viral copies by quantitative PCR

Lung and brain were homogenized in Trizol and RNA was extracted using the Direct-zol RNA Miniprep kit following the manufacturer’s instructions. E gene gRNA was detected using the QuantiNova Probe RT-PCR Kit and protocol and primers (forward primer: 5’-ACAGGTACGTTAATAGTTAATAGCGT-3’, reverse primer: 5’-ATATTGCAGCAGTACGCACACA-3’) and probe (5′-FAM-ACACTAGCCATCCTTACTGCGCTTCG-3IABkFQ-3′) as previously described (86). The standard curve for each PCR run was generated using the inactivated SARS-CoV-2 RNA obtained from BEI (NR-52347) to calculate the viral copy number in the samples. Identical lung and brain portions were utilized for all experiments to generate comparable results.

### Determination of viral titers by TCID_50_ assay

Viral titers from lung and brain homogenate were determined by plating in triplicate on Vero E6 cells (line ATCC CRL-1586 kindly provided by Dr. Sonja Best, NIAID) using 10-fold serial dilutions. Plates were stained with crystal violet after 96 hours to assess cytopathic effect (CPE). Viral titers were determined using the Reed-Muench method.

### Preparation of single cell suspensions from lungs

Lung lobes were diced into small pieces and incubated in RPMI containing 0.33mg/mL Liberase TL and 0.1mg/mL DNase I (both from Sigma Aldrich) at 37°C for 45 minutes under agitation (150rpm). Enzymatic activity was stopped by adding FCS. Digested lung was filtered through a 70μm cell strainer and washed with RPMI. Red blood cells were lysed with the addition of ammonium-chloride-potassium buffer (Gibco) for 3 minutes at room temperature. Cells were then washed with RPMI supplemented with 10% FCS. Live cell numbers were enumerated using AOPI staining on a Cellometer Auto 2000 Cell Counter (Nexcelom).

### Spectral cytometry

To label cells within the pulmonary vasculature for flow cytometric analysis, 2μg anti-CD45 (30-F11; Invitrogen) was administered by intravenous injection 3 minutes prior to euthanasia. Single-cell suspensions prepared from lungs were washed twice with PBS prior to incubating with Live/Dead™ Fixable Blue (ThermoFisher) and Fc Block™ (clone KT1632; BD) for 15 minutes at room temperature. Cocktails of fluorescently conjugated antibodies (listed in **Table 1**) diluted in PBS and 10% Brilliant Stain Buffer (BD) were then added directly to cells and incubated for a further 20 minutes at room temperature. Cells were next incubated in eBioscience™ Transcription Factor Fixation and Permeabilization solution (Invitrogen) for 2-18 hours at 4°C and stained with cocktails of fluorescently labeled antibodies against intracellular antigens diluted in Permeabilization Buffer (Invitrogen) for 30 minutes at 4°C.

**Table 1:**
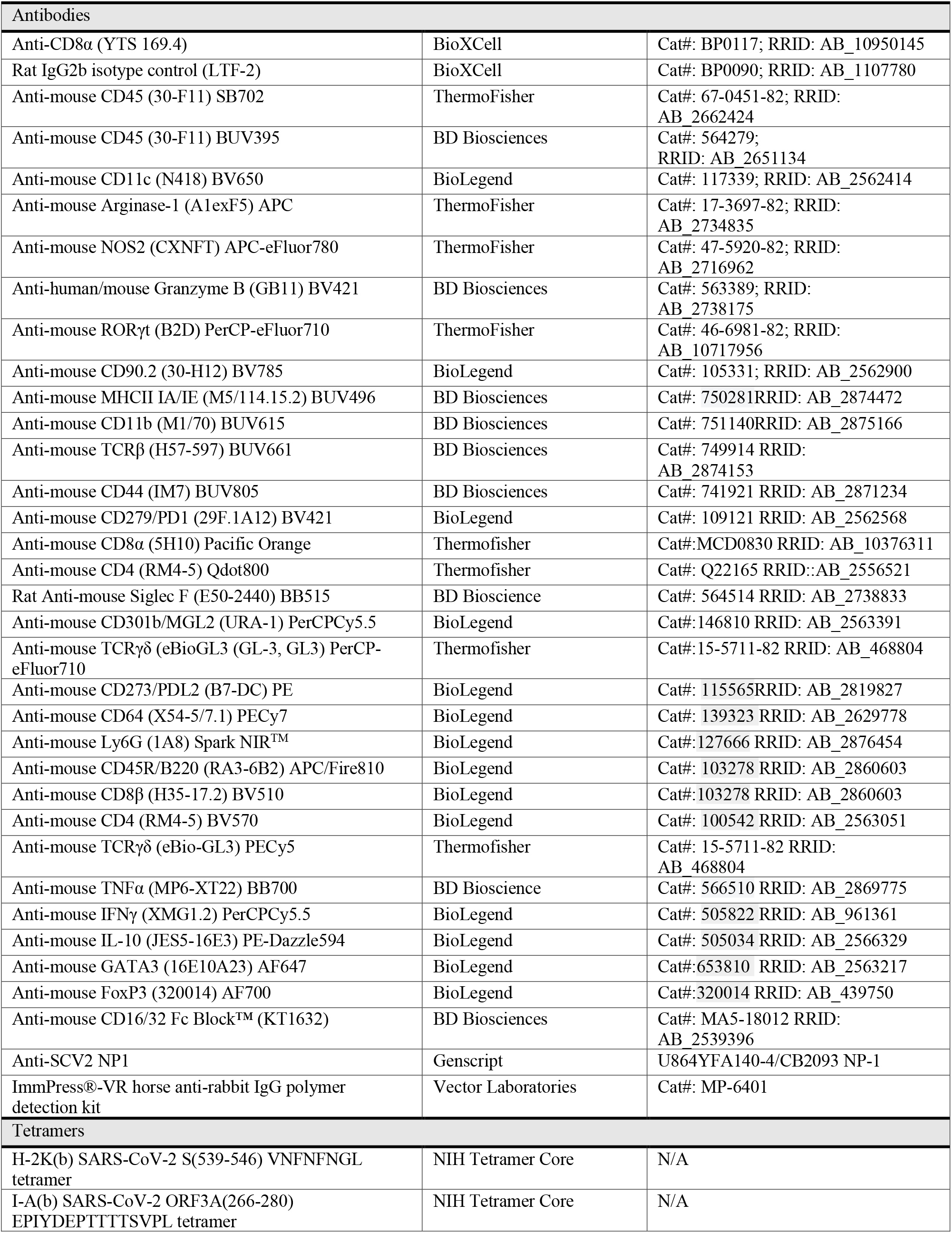
list of antibodies and tetramers.

Spectral Unmixing was performed for each experiment using single-strained controls using UltraComp eBeads™ (Invitrogen). Dead cells and doublets were excluded from analysis. All samples were collected on an Aurora™ spectral cytometer (Cytek) and analyzed using the OMIQ platform (https://www.omiq.ai/) for manual gating of different populations and Joe’s Flow (Github: https://github.com/niaid/JoesFlow) software for unsupervised clustering to identify unique populations in the different groups. Gating strategies are shown in **FigS1**.

### Multiplex Cytokine Array

Cytokines were assessed in lung homogenate using a ProcartaPlex Luminex kit (ThermoFisher) according to the manufacturers’ instructions and measured using a MagPix Instrument (R&D Systems). Total protein was determined by Pierce™ Bradford Assay (ThermoFisher). Cytokine levels were standardized to total protein content.

### Histopathology

Tissues were fixed in 10% neutral buffered formalin for 48-72 hours and then embedded in paraffin. Embedded tissues were sectioned at 5μm and dried overnight at 42°C prior to staining. Specific anti-SCV2 immunoreactivity was detected using a SCV2 nucleoprotein antibody (Genscript) at a 1:1000 dilution. The secondary antibody was the Vector Laboratories ImmPress VR anti-rabbit IgG polymer (cat# MP-6401). The tissues were then processed for immunohistochemistry using the Discovery Ultra automated stainer (Ventana Medical Systems) with a ChromoMap DAB kit (Roche Tissue Diagnostics cat#760– 159). All tissue slides were evaluated by a study-blinded board-certified veterinary pathologist.

### Single cell RNA sequencing

Single cell suspensions were obtained from lungs as described above. Equal number of cells were pooled from all mice in a group. Mice that were found dead or displayed < 5% weight loss in case of infected animals were not pooled. 10,000 cells from each group were loaded on a 10X Genomics Next GEM chip and single-cell GEMs were generated on a 10X Chromium Controller. Subsequent steps to generate cDNA and sequencing libraries were performed following 10X Genomics’ protocol. Libraries were pooled and sequenced using Illumina NovaSeq SP 100 cycle as per 10X sequencing recommendations.

The sequenced data were processed using Cell Ranger (version 6.0) to demultiplex the libraries. The reads were aligned to *Mus musculus* mm10 and SCV2 (MN985325.1) genomes to generate count tables that were further analyzed using Seurat (version 4.1.2). Data are displayed as uniform manifold approximation and projection (UMAP). The different cell subsets from each cluster were defined by the top 50 differentially expressed genes and identification using the SingleR sequencing pipeline (44). Seurat was used for comparisons between each of the different cell cluster of interest either at d0 and at 7dpi. Gene pathway analysis was performed using the publicly available online WebGestalt 2019 analysis toolkit (87) or the Gene ontology Online Resource tool kit (88, 89).

Raw data will be available at publication.

### Visualization

scRNASeq analysis data were visualized using Seurat (version 4.1.2) and R Studio (version 2022.07.1). Cartoons were created using BioRender.com.

### Statistical analysis

Results in graphs are displayed as mean ± SEM using Prism version 7 (GraphPad Software, Inc.) except where mentioned. Statistical analysis was performed using JMP software (SAS, v16) or GraphPad Prism software (v9). Data were analyzed using linear mixed-effects models with a fixed effect of experimental group and a random effect of experiment day. Model assumptions of normality and homogeneous variance were assessed by analysis of the raw data and the model residuals. Right-skewed data were log or square root transformed. In some cases, data were analyzed by Student’s unpaired *t*-test or Mann-Whitney test when comparing two groups, or by One-Way ANOVA with Tukey’s post-test or Kruskal-Wallis test with Dunn’s post-test when comparing three or more groups using GraphPad Prism software (v9). Experimental group was considered statistically significant if the fixed effect F test *p*-value was ≤0.05. Post hoc pairwise comparisons between experimental groups were made using Tukey’s honestly significant difference multiple-comparison test. A difference between experimental groups was taken to be significant if the *p*-value was less than or equal to 0.05 (* *p*<0.05; ** *p*<0.01; *** *p*<0.001; **** *p*<0.0001).

## Author contributions

Conceptualization: KLH, OO, AS, PL. Methodology: KLH, OO, SN. Investigation: KLH, OO, SN, NH, CSC, SDO. Resources: NLG, BAPL, RFJ. Data curation and analysis: OO, KLH, SN, NH, CSC. Writing – original draft: KLH, OO, AS, PL. Writing – review and editing: KLH, OO, CSC, KDM-B, AS, PL. Visualization: OO, KLH, SN, NH, CSC. Supervision: KDM-B, AS, PL. Funding acquisition: AS, PL.

## Acknowledgements

We are grateful to Drs. Paul Baker and Christine Nelson for critical discussion and assistance in setting up SARS-CoV-2 models, the National Cancer Institute Genomics Core for single-cell RNA sequencing; Dr. Craig Martens and the RML Genomics Unit for viral sequencing; the NIAID animal care staff. KLH was partially supported by a Rutherford Postdoctoral Fellowship from Te Apārangi Aotearoa/Royal Society of New Zealand. This research was funded by the Intramural Program of NIAID, NIH.

## Declaration of Interests

The Authors have no conflicts of interest to declare.

---

**Supplementary Figure 1:**
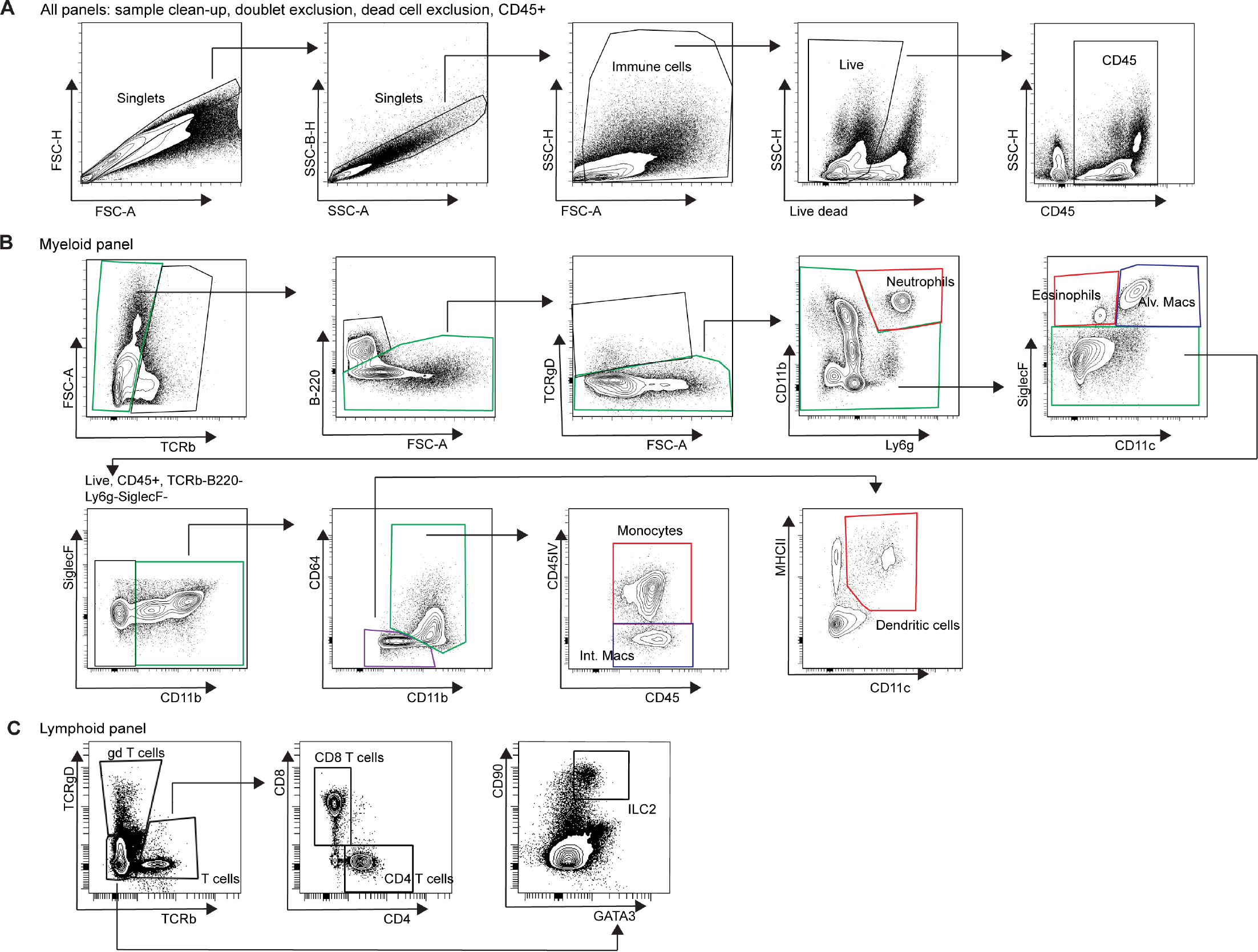
Gating strategies to identify immune cell subsets by spectral cytometry. Single cell suspensions were prepared from the lungs of animals i.v. injected with a fluorescently labeled panCD45 antibody 3 min before euthanasia to allow for identification of cells located within the pulmonary vasculature vs. the cells in the lung interstitium or airways. Strategies employed for sample clean-up (**A**), identifying myeloid populations (**B**) and identifying lymphoid populations (**C**).

**Supplementary Figure 2:**
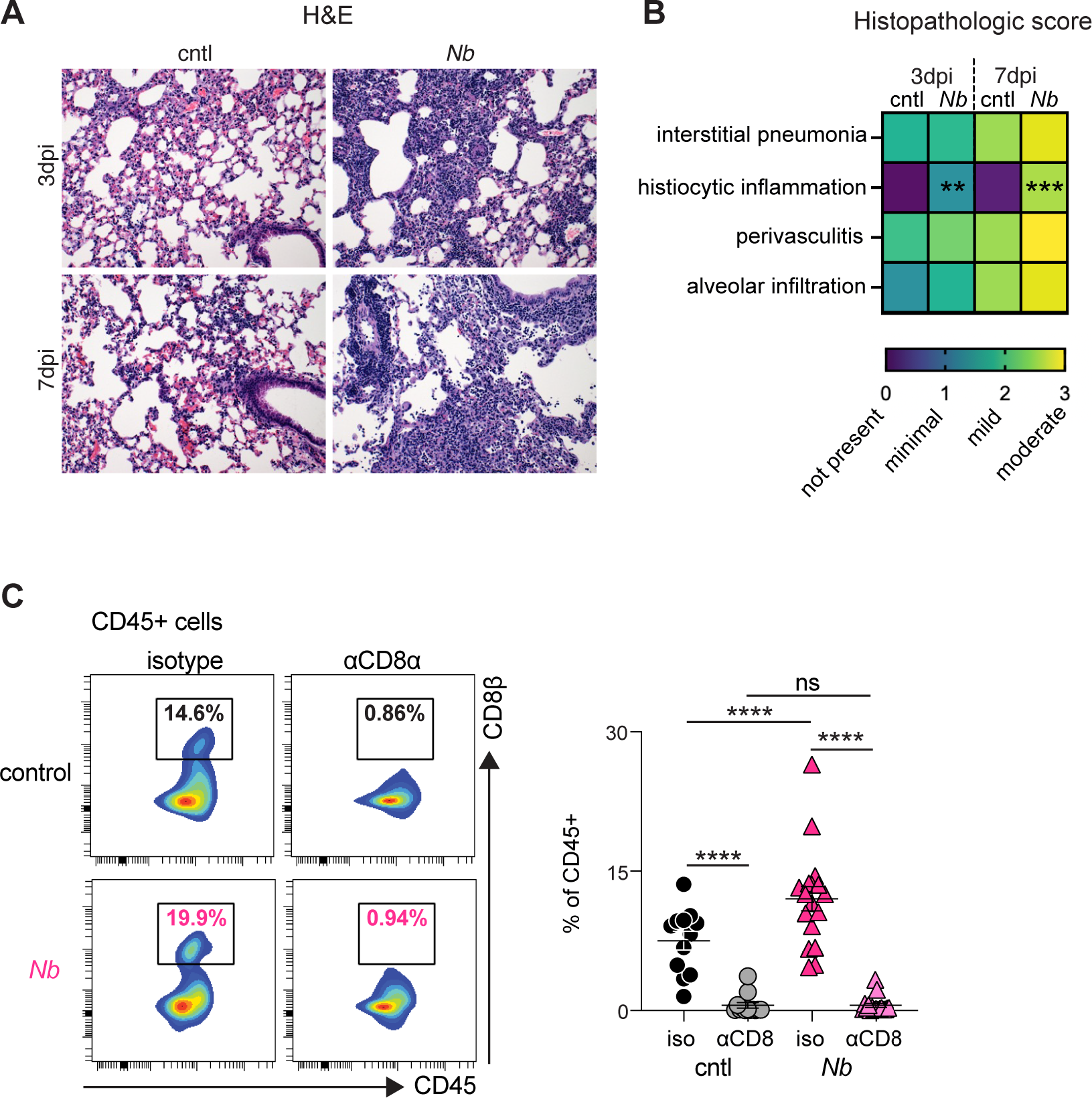
Previous *N. brasiliensis* infection drives lymphocytic inflammation following SCV2 challenge. (**A-B**) K18-hACE2 mice were infected with 500 L3 *N. brasiliensis* (*Nb*) larvae s.c. or left uninfected. After 28 days, animals were challenged i.n. with 10^3^ TCID50 SCV2 and lungs harvested at 3 or 7d post SCV2 for histopathological analysis. *n*=7-10 mice/group; 2 independent experiments. (**A**) Representative hematoxylin and eosin (H&E) stained lung tissue sections. (**B**) Heat map representation of histopathological scores as determined by a board-certified veterinary pathologist. Statistical significance between different groups at different time point was determined using an unpaired Student’s *t*-test with Graph-Pad Prism software. (**C**) K18-hACE2 mice were inoculated with 500 *Nb* larvae by s.c. injection at d-28. Mice were then treated with either anti-CD8α or rat IgG2b isotype control on d-5, d-3, d-1 prior to SCV2 challenge on d0. Lung tissue was harvested at 7dpi to determine frequency of CD8+ T cells by flow cytometry using a CD8β antibody. *n*=11-17 mice/group; 2 independent experiments. Statistical significance was assessed using a linear mixed-effects model with pairwise comparison using JMP software. ns *p*>0.05; * *p*<0.05; ** *p*<0.01; *** *p*<0.001; **** *p*<0.0001

**Supplementary Figure 3:**
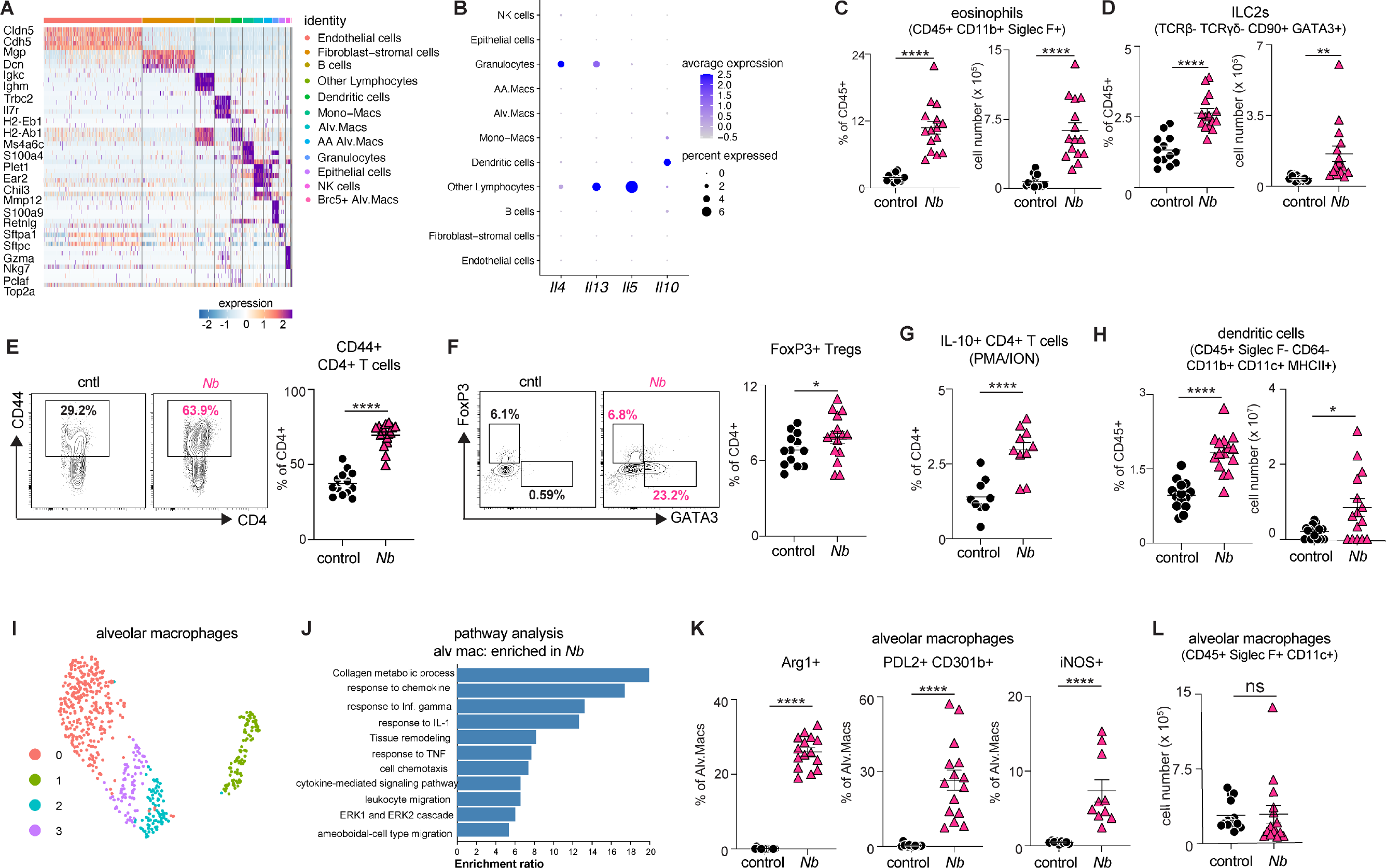
Previous *N. brasiliensis* infection skews the lung micro-environment towards a Type 2 and regulatory phenotype. K18-hACE2 mice were infected with 500 L3 *N. brasiliensis* (*Nb*) larvae s.c. or left uninfected. After 28 days, lungs were harvested and processed for scRNAseq (*n*=pool of 4-5 mice/group), flow cytometric analysis or multiplex cytokine assay (*n*=14-15 mice/group; 3 independent experiments). (**A**) Heat map depicting cluster defining genes used for cell type calling in Fig3A. (**B**) Normalized expression of *Il4*, *Il5*, *Il13* and *Il10* transcripts for each cell type. (**C-K**) Flow cytometric determination of (**C**) the frequency and number of CD11b^+^ Siglec F^+^ eosinophils, (**D**) the frequency and number of TCR^−^ CD90^+^ GATA3^+^ ILC2s, (**E**) frequency of CD44^+^ CD4^+^ T cells, (**F**) frequency of FoxP3^+^ CD4^+^ Tregs, (**G**) frequency of IL-10^+^ CD4^+^ T cells, (**H**) frequency and number of CD64^−^ CD11c^+^ MHCII^+^ CD11b^+^ dendritic cells. Statistical significance was assessed using a linear mixed-effects model with pairwise comparison using JMP software. (**I**) UMAP visualization of Seurat clustering of alveolar macrophages. (**J**) Pathway analysis showing enrichment in *Nb* alveolar macrophages compared to control. (**K**) Flow cytometric assessment of Arg1, PDL2, CD301b and iNOS expression by alveolar macrophages. (**L**) Number of alveolar macrophages as determined by flow cytometry. Statistical significance was assessed using a linear mixed-effects model with pairwise comparison using JMP software. Data are displayed as mean ± SEM. ns *p*>0.05; * *p*<0.05; ** *p*<0.01; *** *p*<0.001; **** *p*<0.0001

**Supplementary Figure 4:**
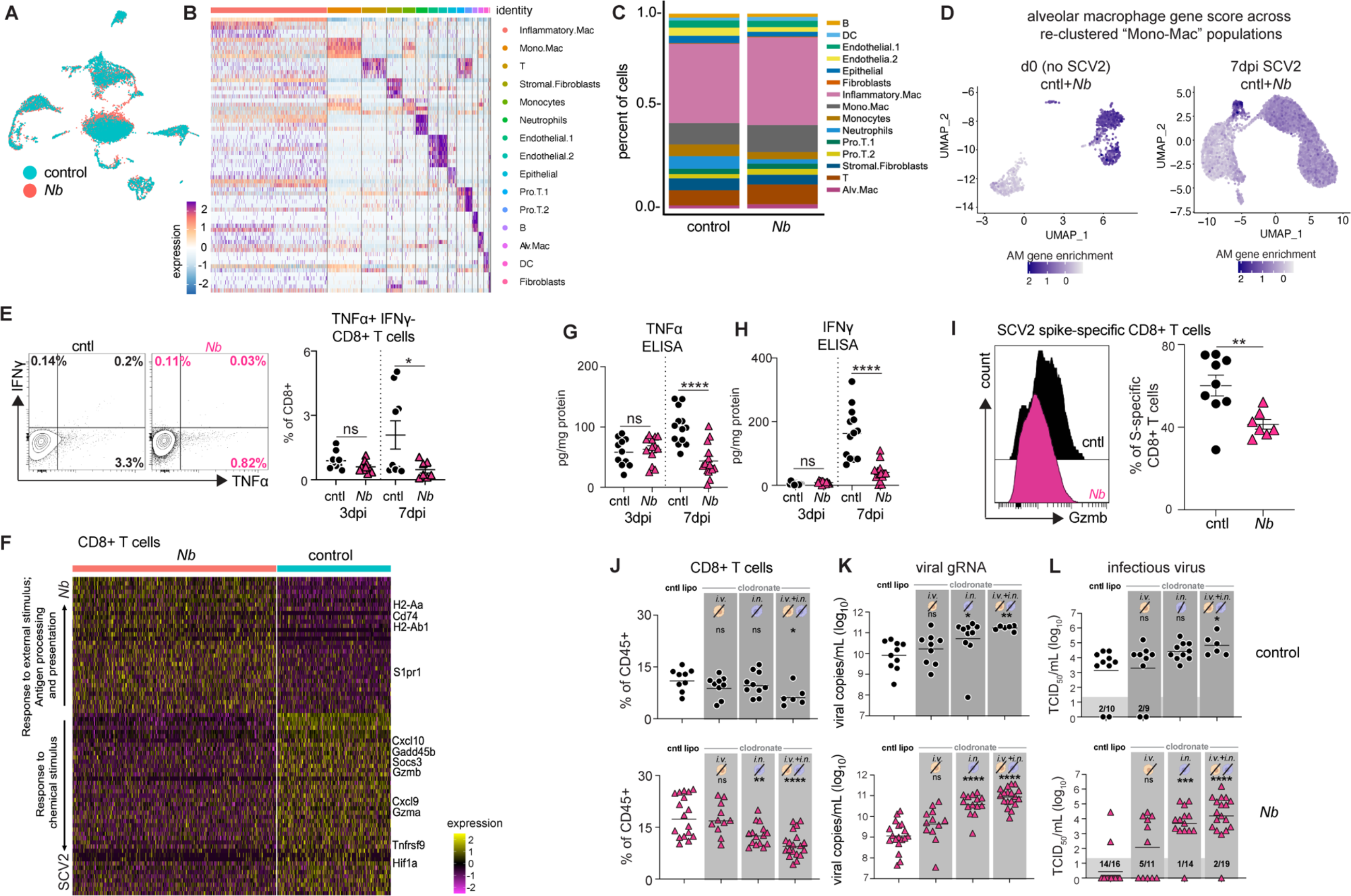
Previous *N. brasiliensis* infection alters the pulmonary macrophage profile and pro-inflammatory cytokine responses after SCV2 challenge. (**A-I**) K18-hACE2 mice were infected with 500 L3 *N. brasiliensis* (*Nb*) larvae s.c. or left uninfected (control). After 28 days, animals were challenged i.n. with 10^3^ TCID50 SCV2. At 3 or 7d post SCV2, lungs were harvested and processed for scRNA-seq (*n*=pool of 3-4 mice/group), flow cytometry or multiplex cytokine assay (*n*=7-15 mice/group; 2-3 independent experiments). Statistical significance was assessed using a linear mixed-effects model with pairwise comparison using JMP software. (**A**) UMAP of all cells separated by experimental group. (**B**) Heat map depicting cluster defining genes used for cell type calling in Fig4A. (**C**) Proportional cellular composition based on scRNA-seq clusters defined in Fig4A. (**D**) Feature plot showing expression of characteristic alveolar macrophage genes, *Marco*, *Mrc1*, *Chil3*, *Car4*, *Ear1*, *Ear2*, *Plet1*, *Fabp1*, *Fabp4* in re-clustered monocyte-macrophage compartment from scRNA-seq data at d0 (Fig3A) and 7dpi SCV2 (Fig4A) (**E**) Frequency of TNFα^+^ CD8^+^ T cells as determined by flow cytometry. (**F**) Differential expression analysis of CD8+ T cells from *Nb* and control samples showing the top 50 DEGs. Enriched pathways are listed on the lefthand side. Each column represents an individual cell. (**G-H**) Protein levels of TNFα (**G**) and IFNγ (**H**) at 3d and 7d post SCV2 as measured in whole lung homogenate by multiplex cytokine assay. (**I**) Granzyme B expression by SCV2 spike-specific CD8+ T cells. (**J-L**) K18-hACE2 mice were inoculated with 500 *Nb* larvae by s.c. injection at d-28. Mice were then treated with either clodronate liposomes or control liposomes by i.n. and/or i.v. administration from d-6 to d-1 prior to SCV2 challenge on d0. Lungs were harvested at 7dpi. *n*=6-19 mice/group; 2 independent experiments. Statistical significance was determined by Kruskal-Wallis test with Dunn’s post-test using the control liposome group as the fixed comparator. (**J**) Frequency of lung CD8+ T cells as determined by flow cytometry, separated by treatment group. Data are mean ± SEM. (**K-L**) Viral loads measured by qPCR (**K**) and TCID50 assay (**L**), separated by treatment group. Geometric mean is shown. Dark gray box indicates values below limit of detection. ns *p*>0.05; * *p*<0.05; ** *p*<0.01; *** *p*<0.001; **** *p*<0.0001

## Notes

### Competing Interest Statement

The authors have declared no competing interest.

## References

1. Contol CfD. 2022. Global cumulative deaths from COVID-19 reported per 100,000 population pp. Global cumulative deaths from COVID-19 reported per 100,000 population https://covid.cdc.gov/covid-data-tracker/#global-counts-rates

2. Mohammadi R, Hosseini-Safa A, Ehsani Ardakani MJ, Rostami-Nejad M. 2015. The relationship between intestinal parasites and some immune-mediated intestinal conditions. Gastroenterol Hepatol Bed Bench 8: 123–31

3. Wolday D, Gebrecherkos T, Arefaine ZG, Kiros YK, Gebreegzabher A, Tasew G, Abdulkader M, Abraha HE, Desta AA, Hailu A, Tollera G, Abdella S, Tesema M, Abate E, Endarge KL, Hundie TG, Miteku FK, Urban BC, Schallig H, Harris VC, de Wit TFR. 2021. Effect of co-infection with intestinal parasites on COVID-19 severity: A prospective observational cohort study. EClinicalMedicine 39: 101054

4. Chacon N, Chacin-Bonilla L, Cesar IM. 2021. Implications of helminth immunomodulation on COVID-19 co-infections. Life Res.

5. Hays R, Pierce D, Giacomin P, Loukas A, Bourke P, McDermott R. 2020. Helminth coinfection and COVID-19: An alternate hypothesis. PLOS Neglected Tropical Diseases 14: e0008628

6. Bradbury RS, Piedrafita D, Greenhill A, Mahanty S. 2020. Will helminth co-infection modulate COVID-19 severity in endemic regions? Nat Rev Immunol 20: 342

7. Ademe M, Girma F. 2021. The Influence of Helminth Immune Regulation on COVID-19 Clinical Outcomes: Is it Beneficial or Detrimental? Infect Drug Resist 14: 4421–6

8. Abdoli A. 2020. Helminths and COVID-19 Co-Infections: A Neglected Critical Challenge. ACS Pharmacol Transl Sci 3: 1039–41

9. Whitehead B, Christiansen S, Østergaard L, Nejsum P. 2022. Helminths and COVID-19 susceptibility, disease progression, and vaccination efficacy. Trends in Parasitology 38: 277–9

10. Osborne LC, Monticelli LA, Nice TJ, Sutherland TE, Siracusa MC, Hepworth MR, Tomov VT, Kobuley D, Tran SV, Bittinger K, Bailey AG, Laughlin AL, Boucher J-L, Wherry EJ, Bushman FD, Allen JE, Virgin HW, Artis D. 2014. Virus-helminth coinfection reveals a microbiota-independent mechanism of immunomodulation. Science 345: 578–82

11. Desai P, Diamond MS, Thackray LB. 2021. Helminth–virus interactions: determinants of coinfection outcomes. Gut Microbes 13: 1961202

12. Rolot M, Dougall AM, Chetty A, Javaux J, Chen T, Xiao X, Machiels B, Selkirk ME, Maizels RM, Hokke C, Denis O, Brombacher F, Vanderplasschen A, Gillet L, Horsnell WGC, Dewals BG. 2018. Helminth-induced IL-4 expands bystander memory CD8+ T cells for early control of viral infection. Nature Communications 9: 4516

13. Furze RC, Hussell T, Selkirk ME. 2006. Amelioration of Influenza-Induced Pathology in Mice by Coinfection with *Trichinella spiralis*. Infection and Immunity 74: 1924–32

14. Scheer S, Krempl C, Kallfass C, Frey S, Jakob T, Mouahid G, Moné H, Schmitt-Gräff A, Staeheli P, Lamers MC. 2014. S. mansoni Bolsters Anti-Viral Immunity in the Murine Respiratory Tract. PLOS ONE 9: e112469

15. McFarlane AJ, McSorley HJ, Davidson DJ, Fitch PM, Errington C, Mackenzie KJ, Gollwitzer ES, Johnston CJC, MacDonald AS, Edwards MR, Harris NL, Marsland BJ, Maizels RM, Schwarze J. 2017. Enteric helminth-induced type I interferon signaling protects against pulmonary virus infection through interaction with the microbiota. Journal of Allergy and Clinical Immunology 140: 1068–78.e6

16. Wescott RB, Todd AC. 1966. Interaction of Nippostrongylus brasiliensis and Influenza Virus in Mice. I. Influence of the Nematode on the Virus. The Journal of Parasitology 52: 242–7

17. Tchuem Tchuenté LA. 2011. Control of soil-transmitted helminths in sub-Saharan Africa: Diagnosis, drug efficacy concerns and challenges. Acta Tropica 120: S4–S11

18. Hotez PJ, Brindley PJ, Bethony JM, King CH, Pearce EJ, Jacobson J. 2008. Helminth infections: the great neglected tropical diseases. J Clin Invest 118: 1311–21

19. Woolhouse MEJ. 1998. Patterns in Parasite Epidemiology: The Peak Shift. Parasitology Today 14: 428–34

20. Harris NL, Loke P. 2017. Recent Advances in Type-2-Cell-Mediated Immunity: Insights from Helminth Infection. Immunity 47: 1024–36

21. Maizels RM, McSorley HJ. 2016. Regulation of the host immune system by helminth parasites. J Allergy Clin Immunol 138: 666–75

22. Colombo SAP, Grencis RK. 2020. Immunity to Soil-Transmitted Helminths: Evidence From the Field and Laboratory Models. Frontiers in Immunology 11

23. Lechner A, Bohnacker S, Esser-von Bieren J. 2021. Macrophage regulation & function in helminth infection. Seminars in Immunology 53: 101526

24. Chen F, Wu W, Millman A, Craft JF, Chen E, Patel N, Boucher JL, Urban JF, Kim CC, Gause WC. 2014. Neutrophils prime a long-lived effector macrophage phenotype that mediates accelerated helminth expulsion. Nature Immunology 15: 938–46

25. Hartung F, Esser-von Bieren J. 2022. Trained immunity in type 2 immune responses. Mucosal Immunology

26. Chen F, El-Naccache DW, Ponessa JJ, Lemenze A, Espinosa V, Wu W, Lothstein K, Jin L, Antao O, Weinstein JS. 2022. Helminth resistance is mediated by differential activation of recruited monocyte-derived alveolar macrophages and arginine depletion. Cell Reports 38: 110215

27. Filbey KJ, Grainger JR, Smith KA, Boon L, van Rooijen N, Harcus Y, Jenkins S, Hewitson JP, Maizels RM. 2014. Innate and adaptive type 2 immune cell responses in genetically controlled resistance to intestinal helminth infection. Immunology and cell biology 92: 436–48

28. Marsland BJ, Kurrer M, Reissmann R, Harris NL, Kopf M. 2008. Nippostrongylus brasiliensis infection leads to the development of emphysema associated with the induction of alternatively activated macrophages. European Journal of Immunology 38: 479–88

29. Reece JJ, Siracusa MC, Southard TL, Brayton CF, Urban JF, Jr., Scott AL. 2008. Hookworm-induced persistent changes to the immunological environment of the lung. Infect Immun 76: 3511–24

30. Lee J-Y, Hamilton SE, Akue AD, Hogquist KA, Jameson SC. 2013. Virtual memory CD8 T cells display unique functional properties. Proceedings of the National Academy of Sciences 110: 13498–503

31. Lanzer KG, Cookenham T, Reiley WW, Blackman MA. 2018. Virtual memory cells make a major contribution to the response of aged influenza-naïve mice to influenza virus infection. Immunity & Ageing 15: 17

32. Arce VM, Costoya JA. 2021. SARS-CoV-2 infection in K18-ACE2 transgenic mice replicates human pulmonary disease in COVID-19. Cell Mol Immunol 18: 513–4

33. Winkler ES, Bailey AL, Kafai NM, Nair S, McCune BT, Yu J, Fox JM, Chen RE, Earnest JT, Keeler SP, Ritter JH, Kang L-I, Dort S, Robichaud A, Head R, Holtzman MJ, Diamond MS. 2020. SARS-CoV-2 infection of human ACE2-transgenic mice causes severe lung inflammation and impaired function. Nature Immunology 21: 1327–35

34. Allen JE, Sutherland TE. 2014. Host protective roles of type 2 immunity: Parasite killing and tissue repair, flip sides of the same coin. Seminars in Immunology 26: 329–40

35. Bouchery T, Volpe B, Shah K, Lebon L, Filbey K, LeGros G, Harris N. 2017. The Study of Host Immune Responses Elicited by the Model Murine Hookworms Nippostrongylus brasiliensis and Heligmosomoides polygyrus. Current Protocols in Mouse Biology 7: 236–86

36. Jiang R-D, Liu M-Q, Chen Y, Shan C, Zhou Y-W, Shen X-R, Li Q, Zhang L, Zhu Y, Si H-R, Wang Q, Min J, Wang X, Zhang W, Li B, Zhang H-J, Baric RS, Zhou P, Yang X-L, Shi Z-L. 2020. Pathogenesis of SARS-CoV-2 in Transgenic Mice Expressing Human Angiotensin-Converting Enzyme 2. Cell 182: 50–8.e8

37. Sun S-H, Chen Q, Gu H-J, Yang G, Wang Y-X, Huang X-Y, Liu S-S, Zhang N-N, Li X-F, Xiong R, Guo Y, Deng Y-Q, Huang W-J, Liu Q, Liu Q-M, Shen Y-L, Zhou Y, Yang X, Zhao T-Y, Fan C-F, Zhou Y-S, Qin C-F, Wang Y-C. 2020. A Mouse Model of SARS-CoV-2 Infection and Pathogenesis. Cell Host & Microbe 28: 124–33.e4

38. Winkler ES, Bailey AL, Kafai NM, Nair S, McCune BT, Yu J, Fox JM, Chen RE, Earnest JT, Keeler SP, Ritter JH, Kang LI, Dort S, Robichaud A, Head R, Holtzman MJ, Diamond MS. 2020. SARS-CoV-2 infection of human ACE2-transgenic mice causes severe lung inflammation and impaired function. Nat Immunol 21: 1327–35

39. Zheng J, Wong LR, Li K, Verma AK, Ortiz ME, Wohlford-Lenane C, Leidinger MR, Knudson CM, Meyerholz DK, McCray PB, Jr., Perlman S. 2021. COVID-19 treatments and pathogenesis including anosmia in K18-hACE2 mice. Nature 589: 603–7

40. Anderson KG, Mayer-Barber K, Sung H, Beura L, James BR, Taylor JJ, Qunaj L, Griffith TS, Vezys V, Barber DL, Masopust D. 2014. Intravascular staining for discrimination of vascular and tissue leukocytes. Nat Protoc 9: 209–22

41. Zhuang Z, Lai X, Sun J, Chen Z, Zhang Z, Dai J, Liu D, Li Y, Li F, Wang Y, Zhu A, Wang J, Yang W, Huang J, Li X, Hu L, Wen L, Zhuo J, Zhang Y, Chen D, Li S, Huang S, Shi Y, Zheng K, Zhong N, Zhao J, Zhou D, Zhao J. 2021. Mapping and role of T cell response in SARS-CoV-2–infected mice. Journal of Experimental Medicine 218

42. Rydyznski Moderbacher C, Ramirez SI, Dan JM, Grifoni A, Hastie KM, Weiskopf D, Belanger S, Abbott RK, Kim C, Choi J, Kato Y, Crotty EG, Kim C, Rawlings SA, Mateus J, Tse LPV, Frazier A, Baric R, Peters B, Greenbaum J, Ollmann Saphire E, Smith DM, Sette A, Crotty S. 2020. Antigen-Specific Adaptive Immunity to SARS-CoV-2 in Acute COVID-19 and Associations with Age and Disease Severity. Cell 183: 996–1012.e19

43. Butler A, Hoffman P, Smibert P, Papalexi E, Satija R. 2018. Integrating single-cell transcriptomic data across different conditions, technologies, and species. Nat Biotechnol 36: 411–20

44. Aran D, Looney AP, Liu L, Wu E, Fong V, Hsu A, Chak S, Naikawadi RP, Wolters PJ, Abate AR, Butte AJ, Bhattacharya M. 2019. Reference-based analysis of lung single-cell sequencing reveals a transitional profibrotic macrophage. Nature Immunology 20: 163–72

45. Nussbaum JC, Van Dyken SJ, von Moltke J, Cheng LE, Mohapatra A, Molofsky AB, Thornton EE, Krummel MF, Chawla A, Liang HE, Locksley RM. 2013. Type 2 innate lymphoid cells control eosinophil homeostasis. Nature 502: 245–8

46. Molofsky AB, Nussbaum JC, Liang HE, Van Dyken SJ, Cheng LE, Mohapatra A, Chawla A, Locksley RM. 2013. Innate lymphoid type 2 cells sustain visceral adipose tissue eosinophils and alternatively activated macrophages. J Exp Med 210: 535–49

47. Van Gool F, Molofsky AB, Morar MM, Rosenzwajg M, Liang HE, Klatzmann D, Locksley RM, Bluestone JA. 2014. Interleukin-5-producing group 2 innate lymphoid cells control eosinophilia induced by interleukin-2 therapy. Blood 124: 3572–6

48. Finkelman FD, Shea-Donohue T, Morris SC, Gildea L, Strait R, Madden KB, Schopf L, Urban JF, Jr. 2004. Interleukin-4- and interleukin-13-mediated host protection against intestinal nematode parasites. Immunol Rev 201: 139–55

49. Nakagomi D, Suzuki K, Meguro K, Hosokawa J, Tamachi T, Takatori H, Suto A, Matsue H, Ohara O, Nakayama T, Shimada S, Nakajima H. 2015. Matrix metalloproteinase 12 is produced by M2 macrophages and plays important roles in the development of contact hypersensitivity. Journal of Allergy and Clinical Immunology 135: 1397–400

50. Nair MG, Parkinson J, Guiliano D, Blaxter M, Allen JE. 2002. IL-4 dependent alternatively-activated macrophages have a distinctive in vivo gene expression phenotype. BMC immunology 3: 1–11

51. Gordon S. 2003. Alternative activation of macrophages. Nature Reviews Immunology 3: 23–35

52. Girgis NM, Gundra UM, Ward LN, Cabrera M, Frevert U, Loke Pn. 2014. Ly6Chigh monocytes become alternatively activated macrophages in schistosome granulomas with help from CD4+ cells. PLoS pathogens 10: e1004080

53. Loke Pn, Allison JP. 2003. PD-L1 and PD-L2 are differentially regulated by Th1 and Th2 cells. Proceedings of the National Academy of Sciences 100: 5336–41

54. Murray PJ, Wynn TA. 2011. Protective and pathogenic functions of macrophage subsets. Nat Rev Immunol 11: 723–37

55. Svedberg FR, Brown SL, Krauss MZ, Campbell L, Sharpe C, Clausen M, Howell GJ, Clark H, Madsen J, Evans CM, Sutherland TE, Ivens AC, Thornton DJ, Grencis RK, Hussell T, Cunoosamy DM, Cook PC, MacDonald AS. 2019. The lung environment controls alveolar macrophage metabolism and responsiveness in type 2 inflammation. Nat Immunol 20: 571–80

56. Bain CC, MacDonald AS. 2022. The impact of the lung environment on macrophage development, activation and function: diversity in the face of adversity. Mucosal Immunol 15: 223–34

57. Del Valle DM, Kim-Schulze S, Huang H-H, Beckmann ND, Nirenberg S, Wang B, Lavin Y, Swartz TH, Madduri D, Stock A, Marron TU, Xie H, Patel M, Tuballes K, Van Oekelen O, Rahman A, Kovatch P, Aberg JA, Schadt E, Jagannath S, Mazumdar M, Charney AW, Firpo-Betancourt A, Mendu DR, Jhang J, Reich D, Sigel K, Cordon-Cardo C, Feldmann M, Parekh S, Merad M, Gnjatic S. 2020. An inflammatory cytokine signature predicts COVID-19 severity and survival. Nature Medicine 26: 1636–43

58. Claassen I, Van Rooijen N, Claassen E. 1990. A new method for removal of mononuclear phagocytes from heterogeneous cell populations in given-names>vitro, using the liposome-mediated macrophage ‘suicide’ technique. Journal of Immunological Methods 134: 153–61

59. Huang L, Nazarova EV, Tan S, Liu Y, Russell DG. 2018. Growth of Mycobacterium tuberculosis in vivo segregates with host macrophage metabolism and ontogeny. Journal of Experimental Medicine 215: 1135–52

60. Vanderheiden A, Thomas J, Soung AL, Davis-Gardner ME, Floyd K, Jin F, Cowan DA, Pellegrini K, Shi PY, Grakoui A, Klein RS, Bosinger SE, Kohlmeier JE, Menachery VD, Suthar MS. 2021. CCR2 Signaling Restricts SARS-CoV-2 Infection. mBio 12: e0274921

61. Lucas C, Wong P, Klein J, Castro TBR, Silva J, Sundaram M, Ellingson MK, Mao T, Oh JE, Israelow B, Takahashi T, Tokuyama M, Lu P, Venkataraman A, Park A, Mohanty S, Wang H, Wyllie AL, Vogels CBF, Earnest R, Lapidus S, Ott IM, Moore AJ, Muenker MC, Fournier JB, Campbell M, Odio CD, Casanovas-Massana A, Herbst R, Shaw AC, Medzhitov R, Schulz WL, Grubaugh ND, Dela Cruz C, Farhadian S, Ko AI, Omer SB, Iwasaki A. 2020. Longitudinal analyses reveal immunological misfiring in severe COVID-19. Nature 584: 463–9

62. Donlan AN, Sutherland TE, Marie C, Preissner S, Bradley BT, Carpenter RM, Sturek JM, Ma JZ, Moreau GB, Donowitz JR, Buck GA, Serrano MG, Burgess SL, Abhyankar MM, Mura C, Bourne PE, Preissner R, Young MK, Lyons GR, Loomba JJ, Ratcliffe SJ, Poulter MD, Mathers AJ, Day AJ, Mann BJ, Allen JE, Petri WA, Jr. 2021. IL-13 is a driver of COVID-19 severity. JCI Insight 6

63. Moran TM, Isobe H, Fernandez-Sesma A, Schulman JL. 1996. Interleukin-4 causes delayed virus clearance in influenza virus-infected mice. J Virol 70: 5230–5

64. Aung S, Tang YW, Graham BS. 1999. Interleukin-4 diminishes CD8(+) respiratory syncytial virus-specific cytotoxic T-lymphocyte activity in vivo. J Virol 73: 8944–9

65. Desai P, Janova H, White JP, Reynoso GV, Hickman HD, Baldridge MT, Urban JF, Jr., Stappenbeck TS, Thackray LB, Diamond MS. 2021. Enteric helminth coinfection enhances host susceptibility to neurotropic flaviviruses via a tuft cell-IL-4 receptor signaling axis. Cell 184: 1214–31.e16

66. Reese TA, Wakeman BS, Choi HS, Hufford MM, Huang SC, Zhang X, Buck MD, Jezewski A, Kambal A, Liu CY, Goel G, Murray PJ, Xavier RJ, Kaplan MH, Renne R, Speck SH, Artyomov MN, Pearce EJ, Virgin HW. 2014. Helminth infection reactivates latent γ-herpesvirus via cytokine competition at a viral promoter. Science 345: 573–7

67. Chetty A, Darby MG, Vornewald PM, Martín-Alonso M, Filz A, Ritter M, McSorley HJ, Masson L, Smith K, Brombacher F, O’Shea MK, Cunningham AF, Ryffel B, Oudhoff MJ, Dewals BG, Layland LE, Horsnell WGC. 2021. Il4ra-independent vaginal eosinophil accumulation following helminth infection exacerbates epithelial ulcerative pathology of HSV-2 infection. Cell Host Microbe 29: 579–93.e5

68. Pedras-Vasconcelos JA, Pearce EJ. 1996. Type 1 CD8+ T cell responses during infection with the helminth Schistosoma mansoni. The Journal of Immunology 157: 3046

69. Chen F, El-Naccache DW, Ponessa JJ, Lemenze A, Espinosa V, Wu W, Lothstein K, Jin L, Antao O, Weinstein JS, Damani-Yokota P, Khanna K, Murray PJ, Rivera A, Siracusa MC, Gause WC. 2022. Helminth resistance is mediated by differential activation of recruited monocyte-derived alveolar macrophages and arginine depletion. Cell Rep 38: 110215

70. Mikhak Z, Strassner JP, Luster AD. 2013. Lung dendritic cells imprint T cell lung homing and promote lung immunity through the chemokine receptor CCR4. Journal of Experimental Medicine 210: 1855–69

71. Yoshie O, Matsushima K. 2015. CCR4 and its ligands: from bench to bedside. International Immunology 27: 11–20

72. Day C, Patel R, Guillen C, Wardlaw AJ. 2009. The chemokine CXCL16 is highly and constitutively expressed by human bronchial epithelial cells. Exp Lung Res 35: 272–83

73. Israelow B, Mao T, Klein J, Song E, Menasche B, Omer SB, Iwasaki A. 2021. Adaptive immune determinants of viral clearance and protection in mouse models of SARS-CoV-2. Science Immunology 6: eabl4509

74. Swain SL, McKinstry KK, Strutt TM. 2012. Expanding roles for CD4+ T cells in immunity to viruses. Nature Reviews Immunology 12: 136–48

75. Chen ST, Park MD, Del Valle DM, Buckup M, Tabachnikova A, Thompson RC, Simons NW, Mouskas K, Lee B, Geanon D, D’Souza D, Dawson T, Marvin R, Nie K, Zhao Z, LeBerichel J, Chang C, Jamal H, Akturk G, Chaddha U, Mathews K, Acquah S, Brown S-A, Reiss M, Harkin T, Feldmann M, Powell CA, Hook JL, Kim-Schulze S, Rahman AH, Brown BD, Beckmann ND, Gnjatic S, Kenigsberg E, Charney AW, Merad M. 2022. A shift in lung macrophage composition is associated with COVID-19 severity and recovery. Science Translational Medicine 14: eabn5168

76. Coakley G, Harris NL. 2020. Interactions between macrophages and helminths. Parasite Immunol 42: e12717

77. Watanabe S, Alexander M, Misharin AV, Budinger GRS. 2019. The role of macrophages in the resolution of inflammation. J Clin Invest 129: 2619–28

78. Spadaro O, Camell CD, Bosurgi L, Nguyen KY, Youm YH, Rothlin CV, Dixit VD. 2017. IGF1 Shapes Macrophage Activation in Response to Immunometabolic Challenge. Cell Rep 19: 225–34

79. Vannella KM, Wynn TA. 2017. Mechanisms of Organ Injury and Repair by Macrophages. Annual Review of Physiology 79: 593–617

80. Chen F, Liu Z, Wu W, Rozo C, Bowdridge S, Millman A, Van Rooijen N, Urban JF, Wynn TA, Gause WC. 2012. An essential role for TH2-type responses in limiting acute tissue damage during experimental helminth infection. Nature Medicine 18: 260–6

81. Wynn TA, Vannella KM. 2016. Macrophages in Tissue Repair, Regeneration, and Fibrosis. Immunity 44: 450–62

82. Hilligan KL, Namasivayam S, Clancy CS, O’Mard D, Oland SD, Robertson SJ, Baker PJ, Castro E, Garza NL, Lafont BAP, Johnson R, Ronchese F, Mayer-Barber KD, Best SM, Sher A. 2021. Intravenous administration of BCG protects mice against lethal SARS-CoV-2 challenge. Journal of Experimental Medicine 219: e20211862

83. Rosas Mejia O, Gloag ES, Li J, Ruane-Foster M, Claeys TA, Farkas D, Wang SH, Farkas L, Xin G, Robinson RT. 2022. Mice infected with Mycobacterium tuberculosis are resistant to acute disease caused by secondary infection with SARS-CoV-2. PLoS Pathog 18: e1010093

84. Hildebrand RE, Chandrasekar SS, Riel M, Touray BJB, Aschenbroich SA, Talaat AM. 2022. Superinfection with SARS-CoV-2 Has Deleterious Effects on Mycobacterium bovis BCG Immunity and Promotes Dissemination of Mycobacterium tuberculosis. Microbiol Spectr 10: e0307522

85. Camberis M, Le Gros G, Urban Jr J. 2003. Animal Model of Nippostrongylus brasiliensis and Heligmosomoides polygyrus. Current Protocols in Immunology 55: 19.2.1–.2.27

86. Corman VM, Landt O, Kaiser M, Molenkamp R, Meijer A, Chu DK, Bleicker T, Brunink S, Schneider J, Schmidt ML, Mulders DG, Haagmans BL, van der Veer B, van den Brink S, Wijsman L, Goderski G, Romette JL, Ellis J, Zambon M, Peiris M, Goossens H, Reusken C, Koopmans MP, Drosten C. 2020. Detection of 2019 novel coronavirus (2019-nCoV) by real-time RT-PCR. Euro Surveill 25

87. Liao Y, Wang J, Jaehnig EJ, Shi Z, Zhang B. 2019. WebGestalt 2019: gene set analysis toolkit with revamped UIs and APIs. Nucleic Acids Research 47: W199–W205

88. Ashburner M, Ball CA, Blake JA, Botstein D, Butler H, Cherry JM, Davis AP, Dolinski K, Dwight SS, Eppig JT, Harris MA, Hill DP, Issel-Tarver L, Kasarskis A, Lewis S, Matese JC, Richardson JE, Ringwald M, Rubin GM, Sherlock G. 2000. Gene Ontology: tool for the unification of biology. Nature Genetics 25: 25–9

8. 2021. The Gene Ontology resource: enriching a GOld mine. Nucleic Acids Res 49: D325–d34

